# Nucleosome positioning stability is a significant modulator of germline mutation rate variation across the human genome

**DOI:** 10.1101/494914

**Authors:** Cai Li, Nicholas M. Luscombe

## Abstract

Nucleosome organization is suggested to affect local mutation rates in a genome. However, the lack of *de novo* mutation and high-resolution nucleosome data have limited investigation. Further, analyses using indirect mutation rate measurements have yielded contradictory and potentially confounded results. Combining >300,000 human *de novo* mutations with high-resolution nucleosome maps, we reveal substantially elevated mutation rates around translationally stable (‘strong’) nucleosomes. Translational stability is an under-appreciated nucleosomal property, with greater impact than better-known factors like occupancy and histone modifications. We show that the mutational mechanisms affected by strong nucleosomes are low-fidelity replication, insufficient mismatch repair and increased double-strand breaks. Strong nucleosomes preferentially locate within young SINE/LINE transposons; subject to increased mutation rates, transposons are then more rapidly inactivated. Depletion of strong nucleosomes in older transposons suggests frequent re-positioning during evolution, thus resolving a debate about selective pressure on nucleosome-positioning. The findings have important implications for human genetics and genome evolution.

## 1 Introduction

Germline *de novo* mutations, which can be passed to offspring, are the primary source of genetic variation in multicellular organisms, contributing substantially to biological diversity and evolution. *De novo* mutations are also thought to play significant roles in early-onset genetic disorders such as intellectual disability, autism spectrum disorder, and developmental diseases (Veltman and Brunner 2012; Acuna-Hidalgo et al. 2016). Thus, investigating the patterns and genesis of *de novo* mutations in the germline is important for understanding genome evolution and human diseases.

Germline and somatic mutation rates vary across the human genome at diverse scales ranging from nucleotide to chromosomal resolution (Hodgkinson and Eyre-Walker 2011; Segurel et al. 2014). Studies revealed factors linked to local mutation rate variation, including sequence context (Michaelson et al. 2012), replication timing (Stamatoyannopoulos et al. 2009), recombination rate (Francioli et al. 2015), DNA accessibility (Sabarinathan et al. 2016) and histone modifications (Michaelson et al. 2012; Schuster-Bockler and Lehner 2012). However, genomic features identified so far explain less than 40% of the observed germline mutation rate variation (at 100Kb to 1Mb resolution) (Terekhanova et al. 2017; Smith et al. 2018). Therefore, important factors remain to be found. Moreover, due to the limited availability of *de novo* mutation datasets, studies focused on coarse-grained mutation rate variation (typically ≥1kb windows for germline data), or used within-species polymorphisms and inter-species divergence whose observations are potentially confounded by natural selection and other evolutionary processes.

Moreover, the underlying mutational processes causing the observed mutation rate variation are poorly understood, though recent studies have highlighted the contributions of error-prone replicative processes (Harris and Nielsen 2014; Lujan et al. 2014; Reijns et al. 2015; Seplyarskiy et al. 2017; Seplyarskiy et al. 2018) and differential DNA repair efficiencies (Supek and Lehner 2015; Perera et al. 2016; Sabarinathan et al. 2016; Frigola et al. 2017). Despite these advances, it remains a challenge to understand the molecular mechanisms associated with mutation rate variation, particularly in the germline.

Here, we focus on the role of nucleosomes in modulating germline mutation rates. Chromatin is considered important because structural constraints could affect the mutability of genomic sequences (Makova and Hardison 2015). Nucleosome organization (including positioning and occupancy) has been reported as a significant factor in humans and other eukaryotes (Sasaki et al. 2009; Tolstorukov et al. 2011; Chen et al. 2012; Michaelson et al. 2012; Lujan et al. 2014; Pich et al. 2018). Studies in different lineages (Sasaki et al. 2009; Tolstorukov et al. 2011; Lujan et al. 2014) reported increased substitution rates around the centers of nucleosomal sequences and increased insertion/deletion rates in linker DNA. However, there are also disagreements between published studies. For example, Michaelson et al. (2012) suggested that high nucleosome occupancy tends to suppress *de novo* mutations, but Smith et al. (2018) found that a comparative analysis using datasets from different studies resulted in opposing conclusions. Due to few available *de novo* mutations for humans, analysis of many studies was based on variant data from within-species polymorphisms or inter-species divergence, which can be affected by natural selection and non-adaptive processes such as GC-biased gene conversion. Furthermore, because of the limitation of available nucleosome maps, some previous studies treated all annotated nucleosomes equally, ignoring the diverse contexts in which they form. Therefore, combined with the scarcity of *de novo* mutation datasets, the effects of nucleosome organization on germline mutation rate variation, particularly at high resolution remain to be elucidated. Here we take advantage of the rapid increase in the number of *de novo* mutation datasets and better understanding of nucleosome organization in the human genome to perform a systematic analysis of this topic.

## 2 Results

### 2.1 Datasets used for analysis

We used >300,000 human *de novo* single-nucleotide variants (SNVs) and >30,000 short insertions/deletions (INDELs), having removed genomic regions that could confound downstream analysis (**Fig. 1a, Supplementary Fig. 1a**; see Methods). Most data come from three large-scale trio sequencing projects which contribute about 100,000 mutations each (Jonsson et al. 2017; Turner et al. 2017a; Yuen et al. 2017). We also examined extremely rare variants (allele frequency ⩽ 0.0001) from the gnomAD database (Lek et al. 2016) which are approximated to *de novo* mutations because they are thought to undergo limited selection and non-adaptive evolutionary processes (Carlson et al. 2018).

**Fig. 1.**
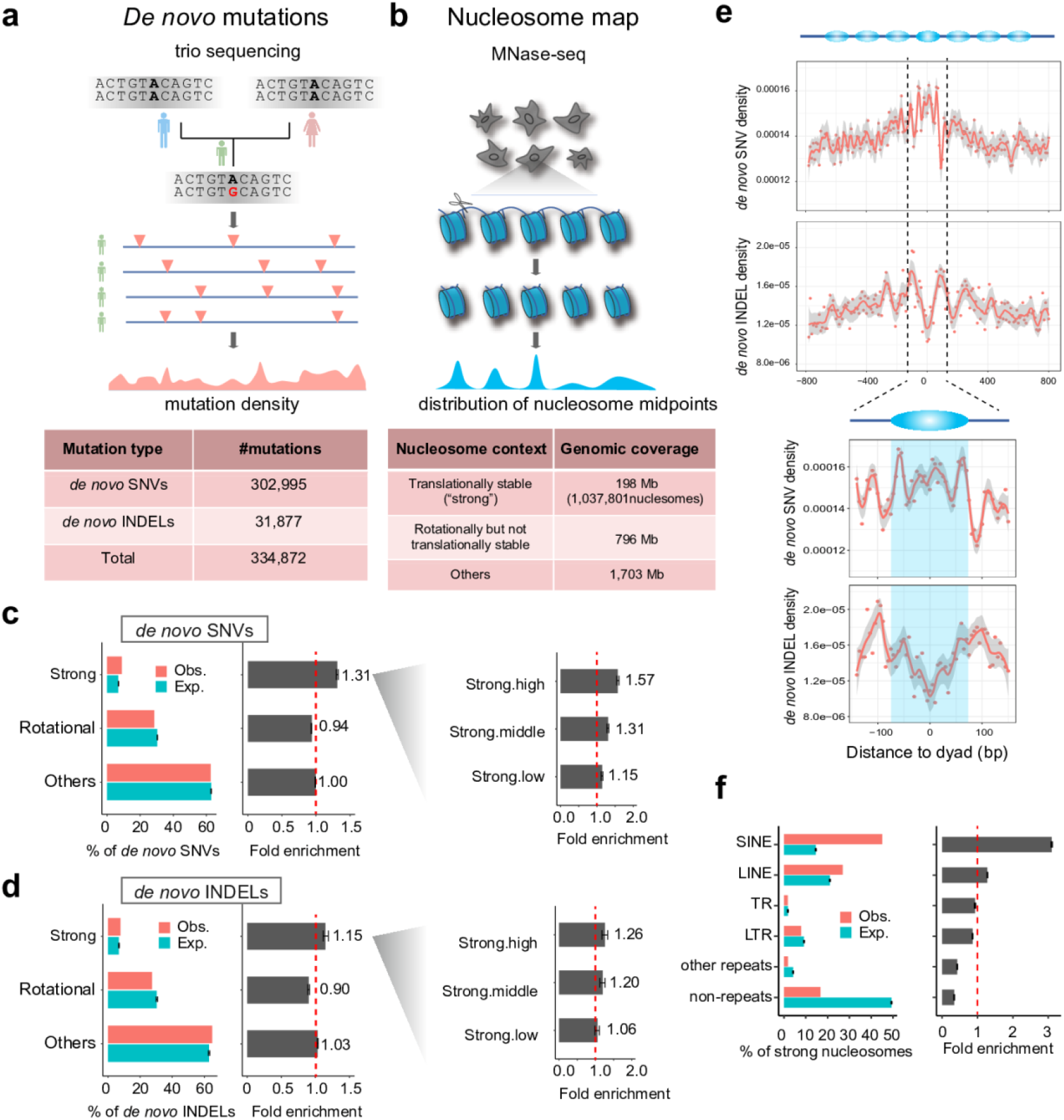
*De novo* mutations are enriched in strong nucleosomes. (**a**) Summary of germline *de novo* mutation data included in study. (**b**) Summary of nucleosome positioning data analysed in study. (**c, d**) Observed versus expected occurrence and fold enrichments of *de novo* (c) SNVs and (d) INDELs in the three different nucleosome contexts. Right-hand panel subdivides strong nucleosomes according to high, medium and low translational stabilities. Error bars depict 95% confidence intervals. (**e**) Top panels, meta-profiles of *de novo* SNV and INDEL densities relative to position of strong nucleosome dyads. Bottom panel, same meta-profiles zoomed into the middle nucleosome. (**f**) Fold enrichment of strong nucleosomes in different repeat elements: SINE (Short Interspersed Nuclear Element), LINE (Long Interspersed Nuclear Element), TR (Tandem Repeat) and LTR (Long Terminal Repeat).

Nucleosome positioning on the genome is described by the translational setting, which defines the location of the nucleosomal midpoint (also called ‘dyad’) and the rotational setting, which defines the orientation of the DNA helix on the histone surface (Gaffney et al. 2012). Using MNase-seq measurements, Gaffney et al. (2012) identified ~1 million ‘strong’ nucleosomes that adopt highly stable translational positioning across seven lymphoblastoid cell lines. Rotationally stable nucleosomes were previously identified from DNase-seq measurements across 43 cell types (Winter et al. 2013), covering 892Mb of the genome. There is a ~50Mb overlap between regions bound by strong nucleosomes and rotationally stable nucleosomes. Using these data, we classified the genome into three groups of regions (**Fig. 1b**; sex chromosomes excluded): i) those containing translationally stable, ‘strong’, nucleosomes (198Mb); ii) those with rotationally but not translationally stable nucleosomes (796Mb); and iii) all other non-N base genomic regions (1,703Mb). West et al. (2014) reported that with the exception of a few specific loci such as transcription start sites, overall nucleosome positioning varies little between cell types. None of the nucleosomal datasets were produced using germ cells, therefore as a precaution we excluded nucleosomes that differ in positioning between cell types (~23Mb; see Methods).

### 2.2 *De novo* SNVs and INDELS are enriched in strong nucleosomes

Genomic regions containing strong nucleosomes have ~30% more *de novo* SNVs (**Fig. 1c**) and ~15% more *de novo* INDELs (**Fig. 1d**) than expected. Similar increases are also apparent for extremely rare variants (**Supplementary Fig. 1b,c**), though effect sizes are smaller than for *de novo* mutations, probably due to the fact that highly mutable sites are under-represented among extremely rare variants (Harpak et al. 2016). Restricting the analysis to strong nucleosomes, we found that those with higher translational stability scores also exhibit higher mutation rates (**Fig. 1c,d**; scores from Gaffney et al., 2012). These results suggest that translational stability is associated with local variation in mutation rates across the genome, a previously unappreciated aspect. Regions containing rotationally stable nucleosomes, in contrast, are slightly depleted of both mutation types; we didn’t perform further analysis on this, as effect of rotational positioning has been comprehensively discussed by Pich et al. (2018). A more detailed view with meta-profiles clearly depicts increased SNV and reduced INDEL densities around dyad regions of strong nucleosomes compared with flanking linker regions (**Fig. 1e**), in line with observations made using polymorphism data (Tolstorukov et al. 2011).

Interestingly, ~80% of strong nucleosomes overlap with repeats (**Fig. 1f, Supplementary Fig. 1d**), especially SINE/Alu (~44%) and LINE/L1 elements (~26%). Genetic variations in repeats are traditionally hard to detect because of poor mappability and so analyses have tended to be cautious in calling variants, resulting in many false negatives (though, few false positives; Lee and Schatz (2012)). Therefore, the above observations probably underestimate the true enrichment of *de novo* mutations in strong nucleosomes. We subdivided strong nucleosomes into three groups: i) Alu-associated, ii) L1-associated and iii) others. Alu-associated nucleosomes display increased SNV rates around the dyads, as seen in the metaprofiles for all strong nucleosomes (**Supplementary Fig. 1e**), whereas non-Alu nucleosomes show increased SNV rates ~60bp away from the dyads, close to the nucleosome edges. Such differences may be due to the different local sequence composition (discussed in next section). In contrast, the patterns of INDEL densities are relatively similar among different groups (**Fig. 1e**).

### 2.3 Controlling for potential confounding factors

Many factors are associated with mutation rate variation. One of the most important is local sequence context - for example, CpG sites are known to be highly mutable and CpG density profiles correlate well with mutation rate profiles in strong nucleosomes (**Supplementary Fig. 1e**). Functional factors like DNA methylation, histone modification, chromatin accessibility, replication timing and recombination rate are also relevant. Therefore, to systematically assess the contribution of nucleosomes to mutation rate variation, we used a logistic regression framework to control for potential confounding factors (**Fig. 2**).

**Fig. 2.**
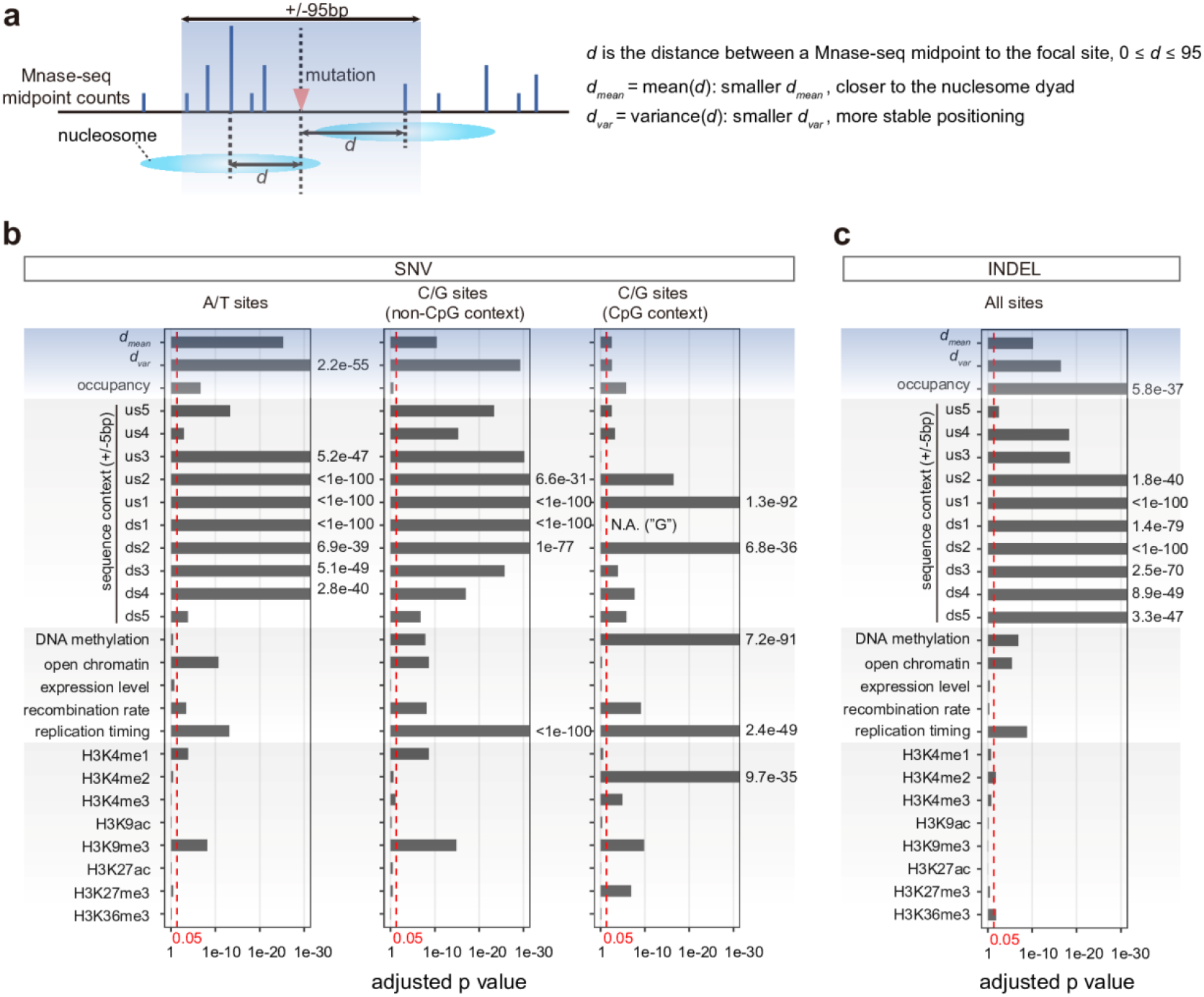
Controlling for potential confounding factors in evaluating contribution of nucleosome organization to mutation rate variation. (**a**) Schematic diagram describing two nucleosome positioning-related variables (***d***_*mean*_ and ***d***_*var*_) relative to a given genomic position. Lower ***d***_*var*_ corresponds to higher translational stability. (**b, c**) Independent statistical significance of potential contributing factors to mutation rate variation, having controlled for other factors; (b) for SNVs and (c) INDELs. Tests for SNVs were performed separately at A/T and C/G sites (non-CpG and CpG contexts respectively). Vertical red lines indicate the threshold for statistical significance (0.05). ‘us’, upstream; ‘ds’, downstream.

We defined three variables to quantify nucleosomal properties relative to a specific nucleotide position in the genome. Two relate to translational positioning: ***d***_*mean*_, the mean distance between the focal position and the midpoints of mapped MNase-seq fragments (maximum distance of 95 bp) and ***d***_*var*_, the variance of these distances (**Fig. 2a**). A smaller ***d***_*mean*_ means that a nucleotide position is closer to nucleosome dyads and a smaller ***d***_*var*_ indicates that the nucleosomes around it are more translationally stable. As the relationship between between ***d***_*mean*_ and SNV rates is non-linear, we defined ***d***_*mean*_ a categorical variable binned into five intervals (Methods; **Fig. 1e, Supplementary Fig. 1e**). The third variable is nucleosome occupancy calculated as a normalised per-base MNase-seq fragment coverage (see Methods). Other factors considered are local nucleotide sequences (±5bp of the focal site) and functional genomic measurements in human germ cells or other cell types if no available germ-cell data (see Methods). ***d***_*var*_ has a relatively weak but statistically significant correlation with many of these factors, suggesting non-independence (**Supplementary Fig. 2**).

To assess the contribution of each factor to local mutation rates, we compared a full logistic regression model encompassing all variables against reduced models missing individual variables; the reported p values indicate how significant a factor is associated with mutation rate variation, having controlled for other factors (**Fig. 2b,c**; Methods). For SNVs, we tested A/T (comprising A>C, A>G and A>T mutations), CpG and non-CpG C/G sites separately (both C>A, C>G and C>T; **Fig. 2b**), whereas they were pooled for INDELs.

Our statistical framework recapitulates reported observations (**Fig. 2b,c, Supplementary Fig. 3**). In agreement with previous studies (Carlson et al. 2018), local sequence context is the biggest contributor to local mutation rate variation (**Fig. 2b,c**), with effect sizes generally declining with increasing distance from the surveyed site. DNA methylation and H3K9me3 are two common epigenetic marks associated with mutation rate variation in general (Schuster-Bockler and Lehner 2012), whereas H3K4me1, H3K4me2, H3K4me3 H3K27me3 and H3K36me3 are linked with specific mutation types. Replication timing has highly statistically significant associations with both SNVs and INDEL mutation types. Recombination rate and open chromatin (measured by ATAC-seq) are also associated with many mutation types. Transcription levels, however, lack any links with local mutation rates here.

Turning to nucleosomal properties, translational stability (***d***_*var*_) is associated with elevated mutation rates at A/T, non-CpG C/G and CpG sites, with the first two showing the greatest effect sizes. INDELs also show similar effects, though the higher p values compared with SNVs could partly be due to the smaller sample size. Examining specific SNV mutation types, ***d***_*var*_ is significantly associated with all A/T and C/G mutations (**Supplementary Fig. 3**), except for CpG>TpG (adjusted p = 0.10).. The regression coefficients for ***d***_*var*_ are always negative (i.e., nucleosome variability is anti-correlated with mutation rate, see coefficients in **Supplementary Table 1**), indicating that translational stability is positively associated with mutation rates thus corroborating the patterns observed in **Fig. 1**. As expected from **Fig. 1**, the mean distance to dyads, ***d***_*mean*_, also displays statistically significant associations with mutations rates at A/T and C/G sites (**Fig. 2b,c**). Finally, nucleosome occupancy is also statistically significant; in contrast to the positioning variables however, here the effect is much larger for INDELs than SNVs (**Fig. 2b,c**; INDELs, adjusted p = 5.8e-37; SNVs, adjusted p = 0.21, 1.6e-6 and 2.2e-7). The regression coefficients of occupancy are negative for SNVs at A/T sites, but positive for SNVs at CpG sites (**Supplementary Table 1**), suggesting that occupancy can have opposing effects on mutability depending on sequence context.

Nucleosome positioning stability is at least partly determined by the occupied DNA sequence and thus its effects on mutation rates to some degree can be attributed to the associated sequence (this also applies to other reported factors such as replication timing). However, higher-order interactions among the long stretches of nucleotides which guide nucleosome positioning are difficult to model properly. Nonetheless, we achieved similar statistical significance for translational stability after including non-additive two-way interaction effects for ±5 nucleotides and the 7-mer mutability estimates from Carlson et al. in regression models (Methods; **Supplementary Fig. 4a,b**).

Since many strong nucleosomes are associated with repeat elements, we added repeat status as a predictor in the regression models (Methods). We still achieved strong statistical significance for translational stability after considering repeat status (**Supplementary Fig. 4c**), suggesting that translational stability is independently associated with mutation rate variation. We also tested repeat and non-repeat regions separately, and in most tests (including those for non-repeat regions) translational stability is a significant factor (**Supplementary Fig. 4d**).

Taken together, the logistic regression modeling analysis recapitulated known factors and confirmed the independent contribution of nucleosome translational stability as a new significant factor to local mutation rate variation.

### 2.4 Mutational processes associated with elevated mutability around strong nucleosomes

#### 2.4.1 Mutational signature analysis

Having established an association between mutation rate and nucleosome translational stability, we next sought to identify mutational mechanisms that might explain it. As an initial screen, we compared the COSMIC mutational signatures for *de novo* mutations within strong nucleosomes and those in genomic background. Mutational signatures were originally developed to infer the mutational processes underlying cancer progression by combining the relative frequencies of 96 possible mutation types (six types of single nucleotide substitutions C>A, C>G, C>T, T>A, T>C and T>G, each considered in the context of the bases immediately 5’ and 3’ to each mutated base; Alexandrov et al. (2013)).

We first consider the relative frequencies of the 96 mutation types in the whole genome and strong nucleosomes in different repeat contexts (**Fig. 3a**). The results account for background differences in trinucleotide frequencies between these regions (Methods). Several mutation types display distinct frequencies in strong nucleosomes, suggesting differences in the underlying mutational processes. For instance, 6 out of 16 T>C mutation types are more prevalent in strong nucleosomes and different repeat-based subgroups display distinct C>T mutation frequencies. L1-associated strong nucleosomes tend to show the most similar mutation frequencies to genomic background, whereas the ‘Others’ group show the most changes, perhaps reflecting the heterogeneity of constituent genomic regions.

**Fig. 3.**
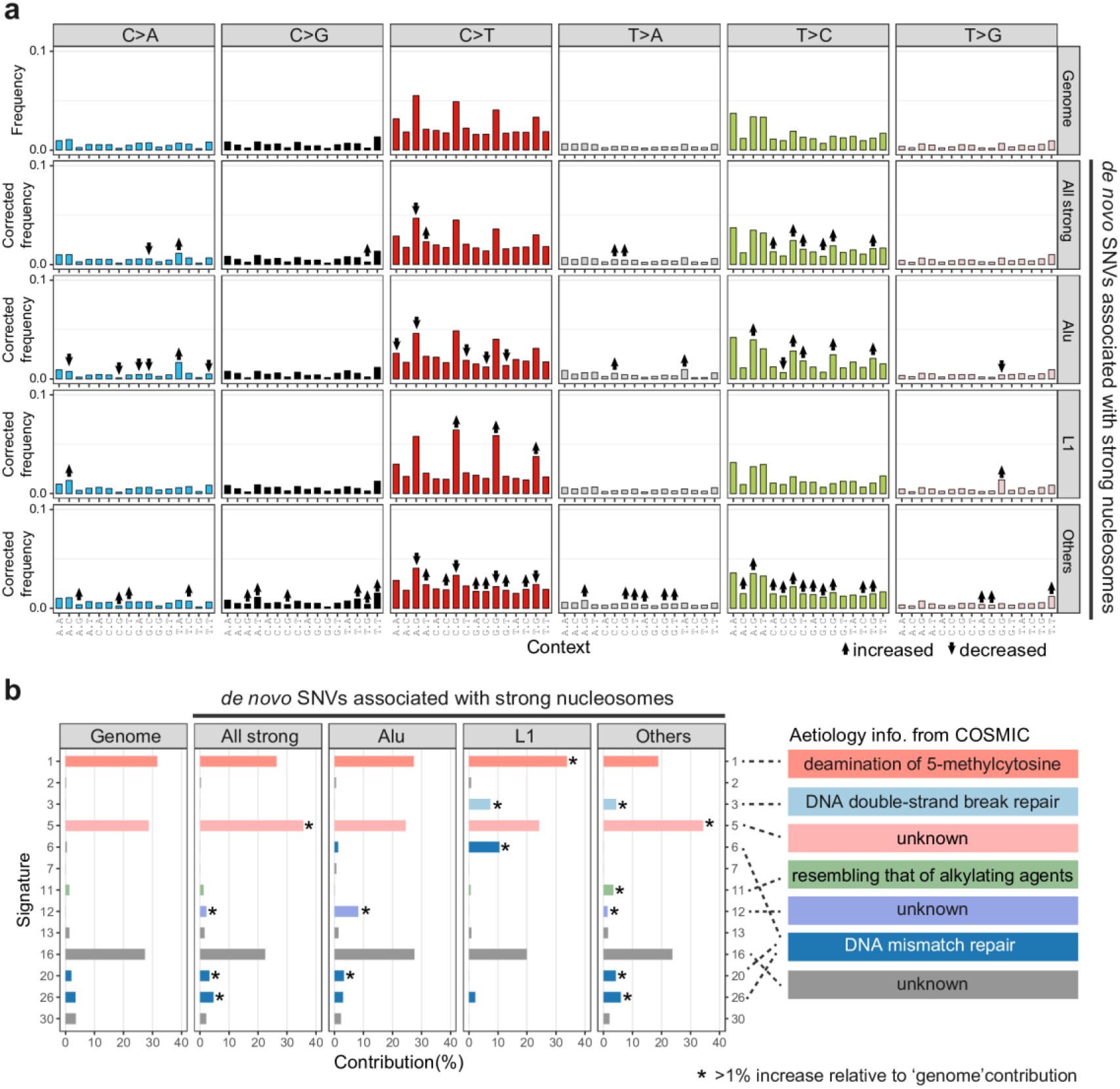
*De novo* SNVs in strong nucleosomes display distinct mutation type frequencies and COSMIC mutational signatures. (**a**) Frequencies of 96 mutation types among *de novo* SNVs; 6 nucleotide substitutions in the context of the bases immediately 5’ and 3’ of the mutated site. SNVs are grouped into those overlapping strong nucleosomes and those elsewhere, and among the former into those overlapping with different classes of repeat elements. ↑ and ↓ indicate mutation types showing statistically significant differences relative to the genomic background SNV set (adjusted p < 0.05, Fisher’s exact test). (**b**) Percentage contribution of COSMIC mutational signatures among different groups of SNVs; only signatures with non-zero values are shown. * indicate mutational signatures displaying >1% increase relative to the genomic background SNV set. Brief summaries of the aetiologies of affected signatures are shown on the right (descriptions taken from the COSMIC website).

Next, we applied the MutationalPatterns software (Blokzijl et al. 2018) to calculate the contribution of COSMIC mutational signatures to different sets of *de novo* SNVs. Three major signatures (Signatures 1, 5 and 16) are present in all tested groups (contributing 87.7% for the whole-genome group, 77.0%~84.5% for strong-nucleosome groups; **Fig. 3b**). Four signatures (Signatures 5, 12, 20 and 26) show increased contribution (>1%) to the ‘all strong-nucleosome’ group relative to the genomic background. The aetiologies of Signatures 5 (~7% increase in strong-nucleosome regions) and 12 (2.2% increase) are currently unknown according to the COSMIC website, but a recent study (Roy et al. 2018) suggested that Signature 5 is likely associated with POL θ-mediated mutagenesis and double-strand break repair. Signatures 20 (1.3% increase) and 26 (1.2% increase) are associated with DNA mismatch repair. There are further differences in associated signatures among strong nucleosome-associated SNVs in different repeat contexts (‘Alu’, ‘L1’ and ‘Others’; **Fig. 3b**), such as signatures 1, 3, 5, 6, 11, 12, 20 and 26. Such differences between different groups could be due to the heterogeneity of contributing mutational processes and redundancy among some COSMIC signatures.

It is worth highlighting that COSMIC mutational signatures were designed for use with cancer genomes and so some germline mutational processes may not be well represented. Nevertheless, our analysis identified several candidate mutational processes associated with strong nucleosomes, such as the mutagenesis linked to DNA mismatch repair (Signatures 6, 20 and 26) and DNA double-strand repair (Signatures 3 and 5). Therefore, to gain deeper insights and to obtain independent evidence for these mutational processes, we examined multiple published genomic and functional genomic datasets below.

#### 2.4.2 Mismatch repair (Signatures 6, 20 and 26)

DNA Mismatch repair (MMR) is a major pathway that is active during DNA replication: it mainly repairs mismatches and short INDELs introduced by DNA synthesis that have escaped polymerase proofreading. Mutations arising from inefficiencies in MMR are represented by Signatures 6, 20 and 26, which show increased contribution to *de novo* SNVs in the ‘All strong nucleosomes’ group (2% increase collectively) and three repeat-based subgroups of mutations (1.6%, 6.7% and 4.3% increase for ‘Alu’, ‘L1’ and ‘Others’, respectively).

We analyzed somatic mutations from two sets of ultra-hypermutated cancer genomes (Campbell et al. 2017). The first comprised genomes with driver mutations in the *POLE* gene encoding the catalytic subunit of DNA polymerase ε (Pol ε, the major replicase for the leading strand) and in one or more of the core MMR genes (*MLH1, MSH2, MSH6, PMS1* and *PMS2*). The second contained cancers with mutated *POLE* but intact MMR. As it is even more challenging to detect somatic mutations in tumor-derived data than re-sequencing of normal individuals, we focused this analysis on strong nucleosomes found in high-mappability regions of the genome (Methods).

We reasoned that differences in mutation distributions between the two sets of genomes could be attributed to the MMR pathway. The overall mutation patterns are similar in both cases, with much higher mutation rates at strong nucleosome boundaries and adjacent linker DNA than the surrounding regions (**Fig. 4a**). This implies that errors introduced during error-prone replication by a deficient Pol ε escape repair by the MMR pathway when they coincide with strong nucleosomes. Next, we calculated an ‘MMR escape ratio’ to quantify the relative amount of replication errors that escapes MMR repair in the *POLE* only mutant cancers compared with the *POLE* and MMR double mutants. Strong nucleosomal regions (especially boundaries and adjacent linkers) display ~10% higher escape ratios than the genome-wide background (**Fig. 4a**). Although A/T sites have higher escape ratios than C/G sites around strong nucleosomes, both C/G and A/T sites exhibit similarly elevated escape ratio profiles, suggesting independence of sequence context.

**Fig. 4.**
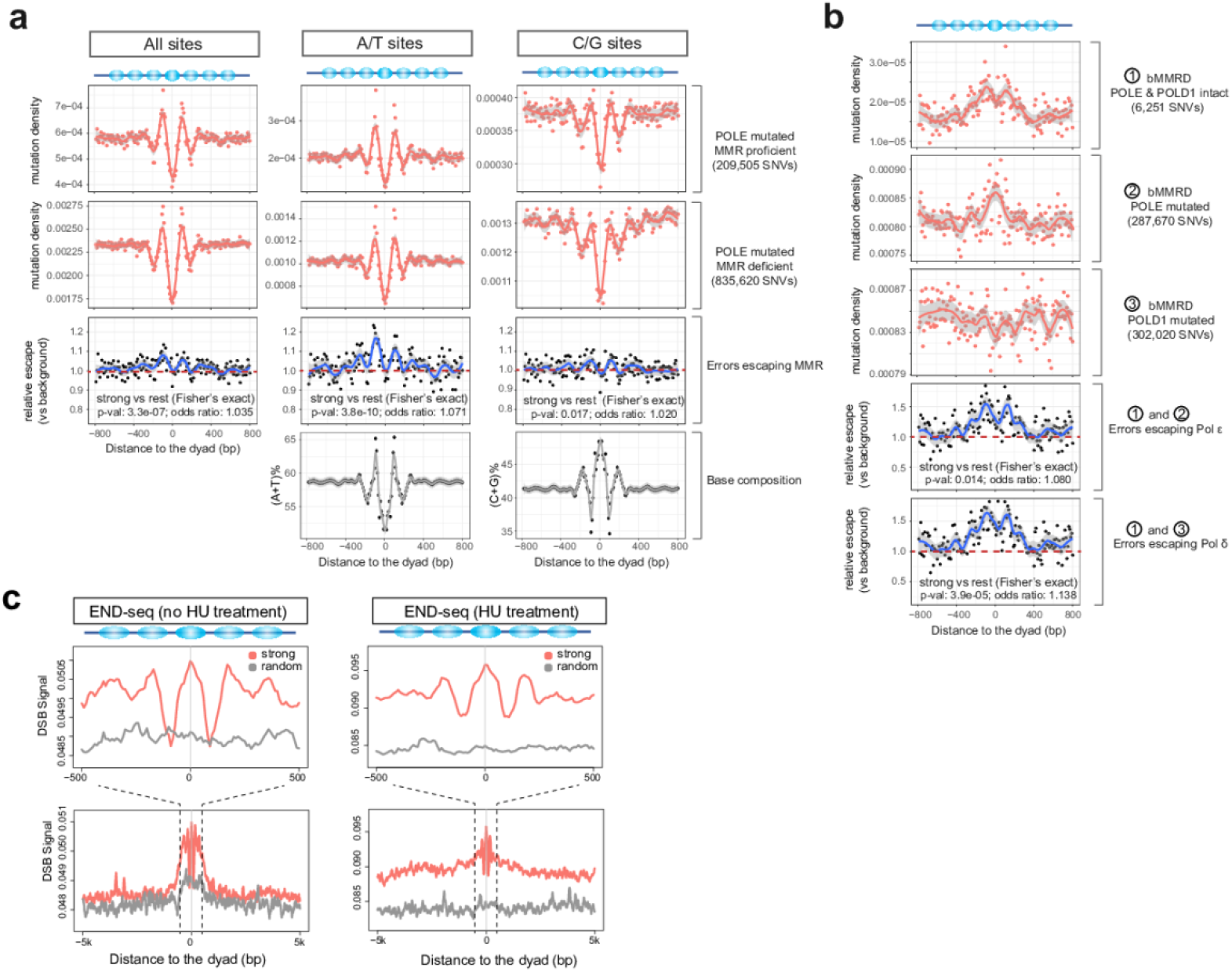
Mismatch repair (MMR), DNA polymerase fidelity and double strand breaks (DSB) explain increased mutation rates in strong nucleosomes. (**a**) Mutation density profiles relative to strong nucleosome dyads in cancer genomes harboring driver mutations in the *POLE* and MMR pathway genes. Numbers of mutations used are indicated in the brackets. The MMR escape ratio compares the mutation densities in the MMR proficient and MMR deficient genomes. (**b**) Mutation density profiles relative to strong nucleosome dyads for bMMRD cancer genomes with different driver mutation statuses in the *POLE* and *POLD1* genes. The escape ratios compare the mutation densities for Pol ε-deficient and Pol δ-deficient cancers with the proficient ones. (**c**) END-seq signal indicating the density of DSBs relative to strong nucleosome dyads. HU, hydroxyurea. Fisher’s exact test was used for testing the association of strong nucleosomal regions (dyad±95bp) with differential MMR/polymerase performance.

Moreover, the apparent ~200-bp periodicity in escape ratio and mutation density profiles are suggestive of associations with nucleosome positioning rather than sequence alone. Together, these observations strongly indicate a relationship between replication errors, MMR and strong nucleosomes in elevating mutation rates.

#### 2.4.3 DNA polymerase fidelity (Signatures 10 and possibly 12)

We also studied the effect of strong nucleosomes on replication fidelity by examining data from children with inherited biallelic mismatch repair deficiency (bMMRD; (Shlien et al. 2015); these include ultra-hypermutated genomes arising from Pol ε and polymerase δ defects (Pol δ, the major replicase for the lagging strand). We estimated Pol δ and Pol ε escape ratios (escaping the proofreading correction of polymerases) using the same reasoning as above (**Fig. 4b**). We found that strong nucleosomes have higher escape ratios for both polymerases relative to the genomic background (**Fig. 4b**), implying that they have lower replication fidelity in these regions. The proofreading escape ratios for both polymerases are even higher than that for MMR (**Fig. 4a,b**) and A/T sites display higher proofreading escape ratios than C/G sites (**Supplementary Fig. 5a**). Again, the periodic pattern in the relative escape profiles (**Fig. 4b, Supplementary Fig. 5a**) suggests that nucleosome positioning contributes to the heterogeneity in replicase fidelity across the genome.

The aetiology of Signature 12 is currently unknown. Here, we found that it contributes 21.15%~21.99% to mutations in POLD1-mutant bMMRD genomes (inferred by MutationalPatterns, **Supplementary Fig. 5b,c**), but much less for other bMMRD samples (0~2.88% for *POLE*-mutant, and 3.32%~10.43% for *POLE/POLD1*-intact). This suggests that Signature 12 is probably associated with Pol δ and that many *de novo* mutations around strong nucleosomes arise from errors escaping Pol δ proofreading. Surprisingly, Signature 10, known to be associated with Pol ε deficiency, is absent from strong nucleosomal *de novo* SNVs (**Fig. 3b**). This suggests that although both Pol ε and Pol δ have high proofreading escape ratios (i.e. low fidelities) around strong nucleosomes (**Fig. 4b**), the majority of the replication errors that are eventually converted to *de novo* mutations are derived from lagging strand replicase Pol δ.

Reijns et al (2015) showed that in budding yeast, Okazaki junctions formed during lagging strand replication tend to be near nucleosome dyads and display elevated mutation rates (Reijns et al. 2015). We tested this by re-analyzing OK-seq data from human lymphoblastoid cells (Petryk et al. 2016). Unlike yeast, Okazaki junctions in humans are more frequently located in the linker regions (**Supplementary Fig. 6**) rather than the dyads, suggesting that the mutagenic effects of Okazaki junctions are different in the two organisms. This may partly be because yeast lacks the typical H1 histone found in human and other eukaryotes. However, the very short reads (single-ended 50bp) of OK-seq data restricted our analysis to nucleosomes with high mappability (~10% of strong nucleosomes), limiting the strength of the conclusions here.

#### 2.4.4 Double-strand breaks (Signatures 3 and 5)

Double-strand break (DSB) repair represented by Signatures 3 and 5 is another potential mechanism involved in strong nucleosome-associated mutations (**Fig. 3b**). Tubbs et al. (2018) studied the genome-wide distribution of DSBs using END-seq and suggested that poly(dA:dT) tracts are recurrent sites of replication-associated DSBs. Our analysis of this data revealed a higher frequency of DSBs around strong nucleosomes compared with genomic background (**Fig. 4c**). The trend holds for experiments with and without hydroxyurea treatment (HU, a replicative stress-inducing agent), suggesting that strong nucleosomes are endogenous hotspots (i.e. without HU treatment) of DSBs during replication. It is notable that young Alu and L1 elements harbor prominent poly(dA:dT) tracts, which are enriched at the boundary and linker regions of strong nucleosomes (**Supplementary Fig. 7a**). The patterns of high DSB frequency still hold true when looking at strong nucleosomes associated with different repeats (**Supplementary Fig. 7b,c**). However, because the END-seq data were sequenced with single-ended 75bp reads and majority of young Alu and L1 elements cannot be assessed with such short reads, we could not pursue further detailed analysis. Since DSB repair can be error-prone (Rodgers and McVey 2016), even using high-fidelity homologous recombination, frequent DSB formation and subsequent error-prone repair likely contribute to the elevated mutation rates around strong nucleosomes.

#### 2.5 Strong nucleosome positioning is mostly associated with young repeat elements and undergoes frequent turnover

Above, we highlighted that ~70% of strong nucleosomes are located in Alu and L1 retrotransposons (**Supplementary Fig. 1d**). Upon examination of the subfamilies (**Fig. 5a,b**), we uncovered a strong enrichment for evolutionarily young L1s (e.g. L1PA2 to L1PA11) and Alus (e.g. AluY to AluSx). Since younger repeats have poorer mappability, these observations probably underestimate the true enrichment. This may also explain why several of the youngest L1 subfamilies (L1PA2 to L1PA5) have lower enrichments than the slightly older subfamilies (**Fig. 5a**).

**Fig. 5.**
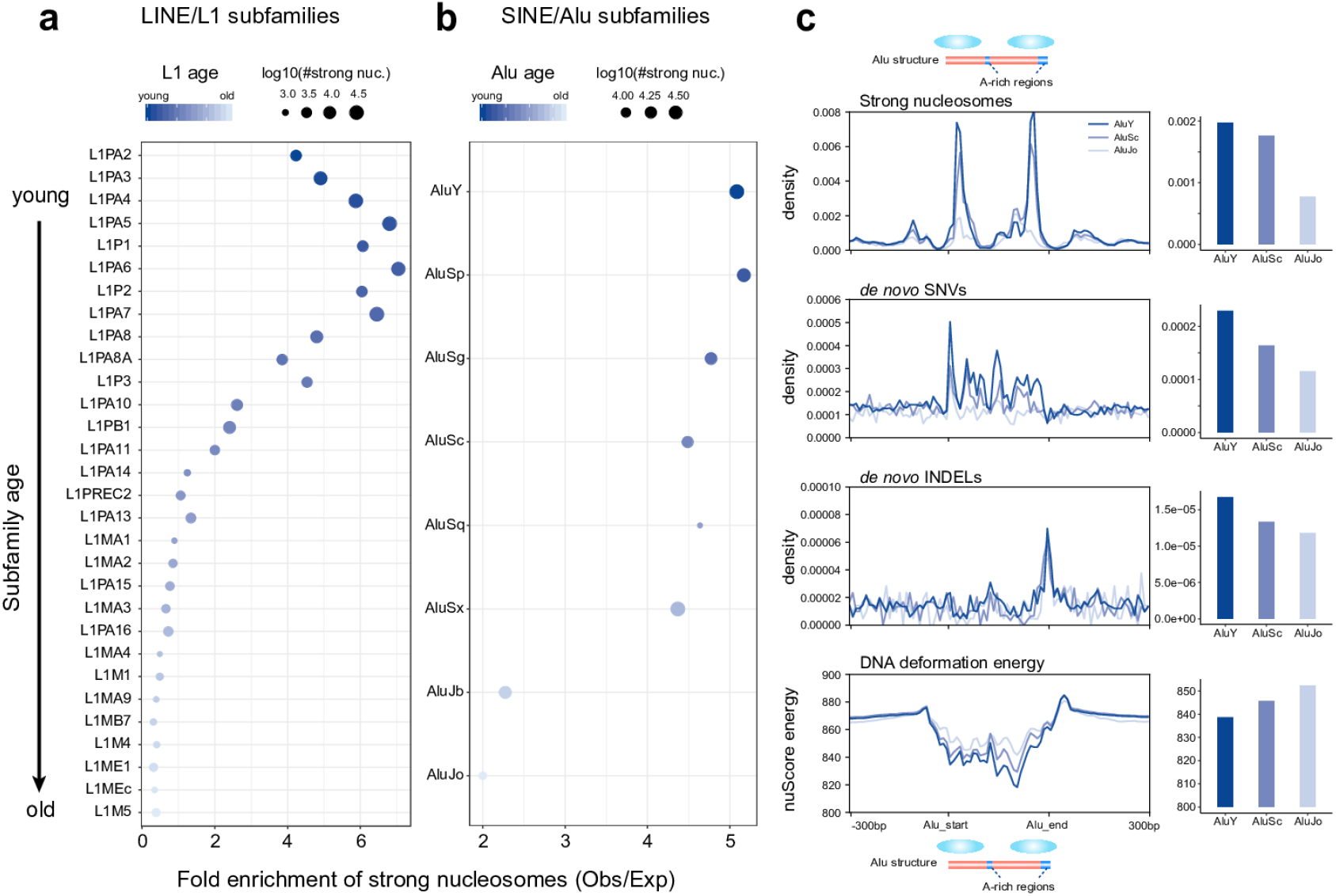
Strong nucleosomes are frequently found inside evolutionarily young LINE and SINE elements. (**a**) Fold enrichment of strong nucleosome occurrence in L1 subfamilies. The top 30 abundant subfamilies are shown ordered by evolutionary age. Dot sizes depict the numbers of strong nucleosomes and color-scale indicates the subfamily age. (**b**) Same as (a) but for Alu elements. (**c**) Densities of strong nucleosome dyads, *de novo* SNVs and *de novo* INDELs along the Alu sequences and flanking regions, grouped by Alu subfamilies of different ages. Bar plots show the average densities for all Alus of different subfamilies on the right. The bottom panel shows the average DNA deformation energies along Alu sequences estimated using nuScore. Profiles were plotted using Alu elements >=250bp and all elements were scaled up to a 300bp region in the plots.

The preference for nucleosomes to occupy specific sections of Alu elements is supported by both *in vitro* and *in vivo* evidence (Englander et al. 1993; Englander and Howard 1995; Salih et al. 2008; Tanaka et al. 2010). We recapitulated these observations for strong nucleosomes using the Gaffney et al. MNase-seq data (**Fig. 5c**): there are two hotspots of strong nucleosomes in young Alus, which fade away in older elements. We also observed that younger Alus exhibit elevated *de novo* mutation rates compared with old ones (**Fig. 5c**), and the weaker translational stability in older Alus is accompanied by reduced *de novo* mutation rates for both SNVs and INDELs (**Fig. 5c**). Thus, there is an intriguing interplay between Alus, strong nucleosomes and mutation rates.

The histone octamer is thought to preferentially bind DNA sequences presenting lower deformation energy costs (Tolstorukov et al. 2008). We estimated deformation energies using the nuScore software (Tolstorukov et al. 2008) based on the DNA sequence and nucleosome core particle structure and we found that Alus do indeed exhibit lower deformation energies than surrounding regions (**Fig. 5c**). Furthermore, the energies of Alu elements tend to increase with age, suggesting that the accumulated mutations in Alu sequences reduced their nucleosome-binding stability. This is also supported by comparing deformation energies of Alu consensus sequences (ancestral states) and those of current genomic sequences (**Supplementary Fig. 8a**). We further analyzed the 3’ end sequences of L1 elements harboring strong nucleosomes and observed similar patterns (**Supplementary Fig. 8b,c**).

Studies have suggested that natural selection appears to preserve nucleosome positioning during evolution (Prendergast and Semple 2011; Tolstorukov et al. 2011; Drillon et al. 2016), but they had differing views about the effects of selection on the underlying sequence. In contrast, Warnecke et al. (2013) suggested that the observed sequence divergence patterns around nucleosomes can be explained by frequent nucleosome re-positioning after mutation, rather than by natural selection. Since these results were mainly based on human polymorphisms or inter-species divergence, indirect mutation rate measurements were potentially confounded by selection and non-adaptive processes. The use of *de novo* mutations helps resolve this debate to some extent.

As we showed above, there is considerable *de novo* mutation rate variation around strong nucleosomes (**Fig. 1e, Supplementary Fig. 1**), which cannot be ignored in any selection analysis. Furthermore, strong nucleosomes are clearly preferentially present in young SINE/LINE elements and the strength of translational stability decays substantially over time (**Fig. 5**). These observations support the re-positioning model over a long evolutionary scale. Since a large majority of strong nucleosomes associated with SINE/LINE elements are expected to become non-strong ones in future, selection for preserving positioning might not be as widespread as previously suggested, though it may happen at some particular regions or within a short evolutionary scale.

### 3 Discussion

Though the involvement of nucleosome organization in DNA damage/repair processes was recognised nearly 30 years ago (Smerdon 1991), its genome-wide effects on germline mutation rates (particularly in higher eukaryotes) have remained poorly understood. Our analysis combining large-scale *de novo* mutation and nucleosome datasets in human provides several important insights into this topic.

A major finding is that strong translational positioning of nucleosomes is associated with elevated *de novo* mutation rates, which is also supported by observations using extremely rare variants in polymorphism data. The ability to use *de novo* mutations here allowed us to bypass confounding evolutionary factors such as selection, thus allowing direct assessment of the impact on background mutation rates. Importantly, our statistical tests controlling for nucleosome occupancy and other related factors confirmed the significant contribution of translational stability to mutation rate variation. Therefore, we have discovered a novel factor that significantly modulate germline mutation rate variation.

Investigating the underlying mutational processes responsible for this association remains challenging. Nevertheless, we obtained several informative results regarding potential mechanisms by leveraging published omics data related to DNA damage and repair. In doing so, we revealed that MMR, replicase fidelity and DSB contribute significantly to elevated mutation rates around strong nucleosomes. In particular, multiple sets of ultra-hypermutated cancer data allowed us to quantify the performance of MMR and replicases by calculating the repair escape ratios. The results probably apply to germ cells because i) they agree nicely with the observations from our mutational signature analysis with *de novo* mutations and ii) recent studies suggested that replicative errors account for majority of mutations arising in both somatic and germ cells (Tomasetti and Vogelstein 2015; Tomasetti et al. 2017). The precise molecular interactions determining the relationships between strong nucleosome positioning, replicase fidelity and DNA repair are still not clear. However, based on the evidence from our analysis with the omics data and previous studies (Li et al. 2009; Reijns et al. 2015; Tubbs et al. 2018), we speculate that strong nucleosomes may act as particularly strong barriers which impair the performance of the replication and repair machineries. There may be additional, unexamined effects on DNA damage/repair processes related to germline development, but many published genomic datasets about DNA damage/repair were generated in non-germ cells and with very short sequencing reads (e.g. <100bp), which hinder accurate analysis. Improved sequencing strategies such as long-read sequencing and direct measurement in germ cells would benefit future related studies.

Interestingly, we found that strong nucleosomes are preferentially located within young LINE and SINE elements, two of the most common retrotransposons in the human and other mammalian genomes. Owing to their potentially deleterious effects, newly inserted retrotransposons are tightly repressed by multiple regulatory mechanisms, such as DNA methylation and H3K9me3 (Slotkin and Martienssen 2007). Strong nucleosome positioning, which may mask access to the transcription machinery, could be another layer of the repressive system. Furthermore, the hypermutation in young SINEs/LINEs, partly contributed by associated strong nucleosomes, could lead to the rapid reduction of retrotransposition capacity. Therefore, the combination of strong nucleosome positioning and hypermutation in SINEs/LINEs might have facilitated their expansion across the genome.

The decreasing numbers of strong nucleosomes in older LINE/SINE elements imply widespread nucleosome re-positioning during evolution. Since nucleosome positioning is strongly affected by the underlying DNA sequence, their re-positioning probably arises from the accumulation of mutations. Our data largely disagree with the previous hypothesis of widespread selection for maintaining nucleosome positioning in the human genome (Prendergast and Semple 2011). Another reason for favoring the re-positioning model is that most genomic regions do not employ strong positioning, possibly due to its relatively high mutagenic potential.

Finally, we summarized our major findings in a proposed model in **Fig. 6**, which demonstrates the relationship among nucleosome positioning, mutation rate variation, retrotransposons and evolution. Given the importance of germline *de novo* mutations in evolution and human diseases and the universal roles of nucleosomes in eukaryotic genome organization and regulation, our work should have profound implications in related research areas.

**Fig. 6.**
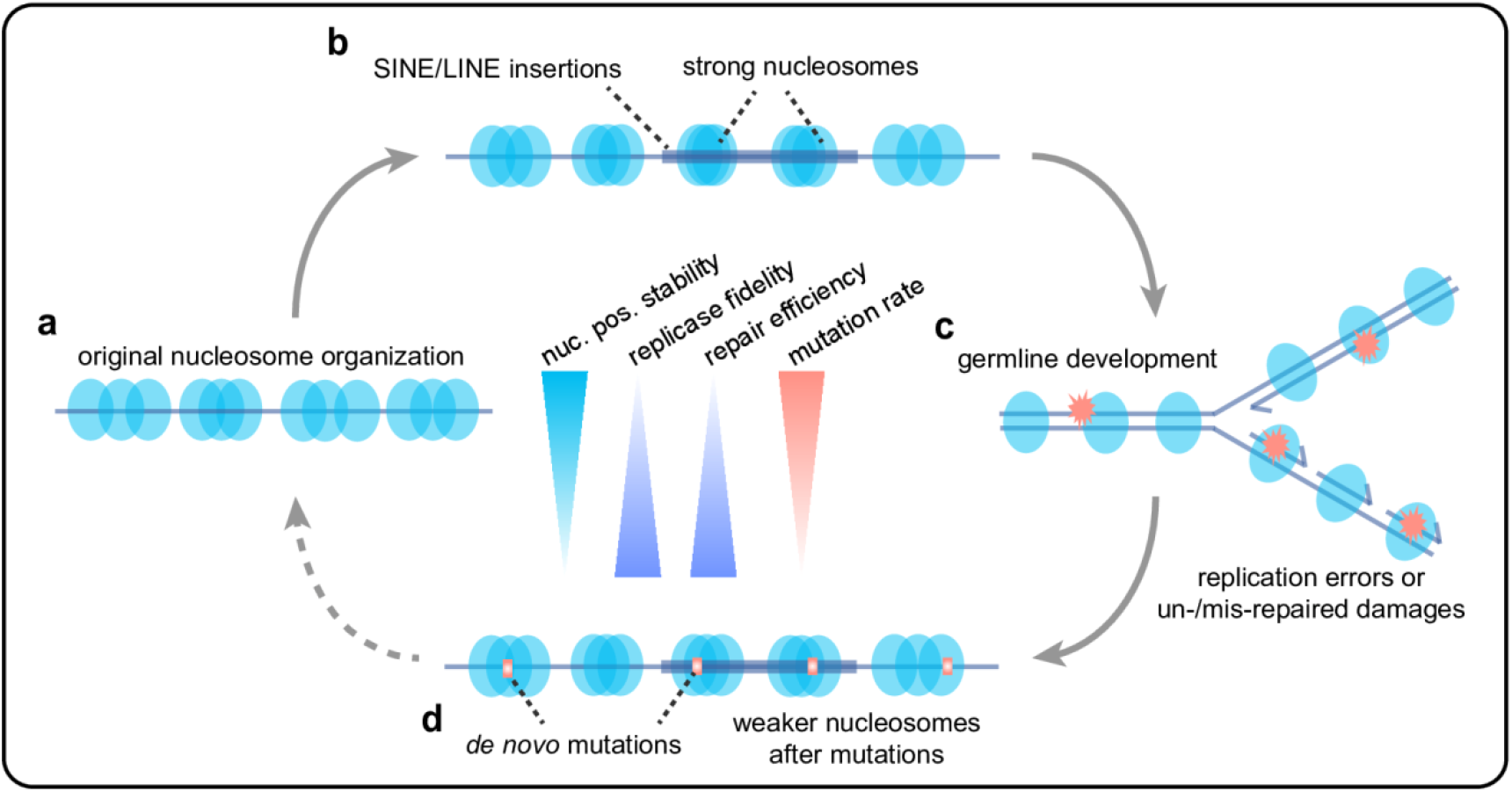
Proposed model of the interplay between nucleosome translational stability, mutation rate and transposable elements. (**a**) Most genomic regions are occupied by nucleosomes lacking strong translational stability. (**b**) Strong nucleosomes are preferentially associated with newly inserted SINE/LINE elements. (**c**) Strong nucleosomal regions are subject to high mutation rates during germline development, caused by mutational processes such as low replicase fidelity, inefficient MMR and DSB repair. (**d**) Accumulation of mutations reduces translational stability of strong nucleosomes and reduces transposition capacity of transposable elements.

## Methods

### Mutation datasets

*De novo* mutations identified in multiple large-scale trio sequencing project were downloaded from denovo-db v1.6.1 (Turner et al. 2017b). Seven studies with >1000 *de novo* mutations (Genome of the Netherlands 2014; Turner et al. 2016; Yuen et al. 2016; Jonsson et al. 2017; Turner et al. 2017a; Yuen et al. 2017; Werling et al. 2018) were considered in our analysis (**Supplementary Fig. 1a**). Extremely rare variants (derived allele frequency ≤ 0.0001) were obtained from Genome Aggregation Database (gnomAD, release 2.0.2) (Lek et al. 2016).

### Nucleosome datasets

We used the 1,037,801 strong nucleosomes (i.e. translationally stable nucleosomes) identified based on MNase-seq data of sequenced seven lymphoblastoid cell lines from Gaffney et al. (Gaffney et al. 2012). The original hg18-based coordinates of annotated nucleosomes were converted to hg19 using the ‘liftOver’ tool from UCSC genome browser. The rotationally stable nucleosomes identified based on 49 DNase-seq samples (43 distinct cell types) were from Winter et al. (Winter et al. 2013). We classified the human genome into three groups based on the nucleosome contexts (**Fig. 1b**): i) regions covered by translationally stable (‘strong’) nucleosomes; ii) regions covered by rotationally but not stable translationally nucleosomes; and iii) the remaining genomic regions. Chromosomes X and Y were excluded from analysis as some other datasets used in our work lacked data for these chromosomes. As the nucleosome maps we used were not derived from germ cells, for downstream analysis we excluded the genomic regions in which nucleosome positioning were found to differ between human embryonic stem cells and differentiated fibroblasts (West et al. 2014). Based on the positioning stability scores defined in Gaffney et al., we divided the one million strong nucleosomes into three categories of equal sizes with different levels of stability – ‘high’, ‘middle’ and ‘low’, which were used for analysis shown in **Fig. 1** and **Supplementary Fig. 1.**

### Accounting for mappability

Sequencing read mappability can significantly affect variant calling results and other aligned read-depth based measurements (e.g. nucleosome occupancy). The sequencing reads for detecting *de novo* mutations used in our analysis were mainly 150bp paired-end reads, with fragment sizes ranging from 300-700bp (**Supplementary Fig. 1**). We used the Genome Mappability Analyzer (GMA) (Lee and Schatz 2012) to generate the mappability scores for simulated paired-end 150 reads with fragment sizes set to be 400bp. Only the regions with GMA mappability scores of >=90 (~2.59Gb) were considered for most analyses, unless specified otherwise. We did not use the mappability tracks from ENCODE for the *de novo* mutation data, because those tracks were only for single-ended reads. For some analyses, additional filtering were applied if other associated datasets suffered from more severe mappability issues. For measuring nucleosome occupancy, we used the method described in the Gaffney et al. to simulate paired-end 25bp reads matching the base compositions of MNase-seq data in the human genome, and then calculated per-base coverage depth by the simulated fragments. The 10bp-bin ratios between the MNase-seq read coverage and the simulated read coverage were used for measuring the occupancy.

### Enrichment analysis for *de novo* mutations in different nucleosome contexts

Genomic association tester (GAT) (Heger et al. 2013), a tool for computing the significance of overlap between multiple sets of genomic intervals, was used to estimate the expected numbers of mutations in different contexts (sampling >=1000 times), which were then compared with the observed numbers. Low-mappability regions were excluded from analysis. A similar analysis was also done for the extremely rare variants of gnomAD. Analysis of meta-profiles along strong nucleosomes was done using deepTools (Ramirez et al. 2014).

### Statistical modelling of the contribution of different factors to mutation rate variation

As described in the main text, for a given genomic position, we defined two variables regarding the translational positioning of nearby nucleosomes (**Fig. 2a**):

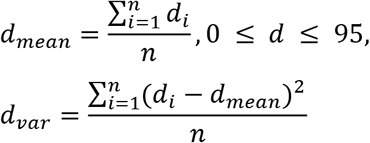

where *d* is the distance between a MNase-seq midpoint to the focal site. We considered MNase-seq midpoints within ±95bp of the focal site, because genome-wide nucleosome repeat length was estimated to be 191.4bp for the Gaffney et al. data (Gaffney et al. 2012). Genomic sites without any MNase-seq midpoint within ±95bp were excluded from analysis (123Mb out of 2.59Gb excluded). The measurements for nucleosome occupancy were 10bp-bin ratios between the MNase-seq read coverage and the simulated read coverage. We did not use the positioning score *S(i)* defined in Gaffney et al. to measure positioning stability in our modelling analysis, because *S(i)* was designed for identifying the stable dyads and so for non-dyad positions it does not represent the positioning stability properly.

RNA expression, DNA methylation and chromatin accessibility (ATAC-seq) data from human spermatogonial stem cells were from Guo et al. (Guo et al. 2017). For the RNA-seq and ATAC-seq data from Guo et al., because the genome-wide read signal tracks were not available, we downloaded, processed and mapped the raw reads to generate the genome-wide tracks. Since suitable data for histone modifications in human germ cells were not available, we used the ChIP-seq data of human embryonic stem cells from ENCODE (ENCODE Consortium 2012). Replication timing data (Repli-seq of GM12878) were also from ENCODE. The data of recombination rates were from the HapMap project (International HapMap Consortium et al. 2007).

A binary logistic regression framework was used to assess the contribution of different factors to mutation rate variation across the genome systematically. The logistic regression model is described as below:

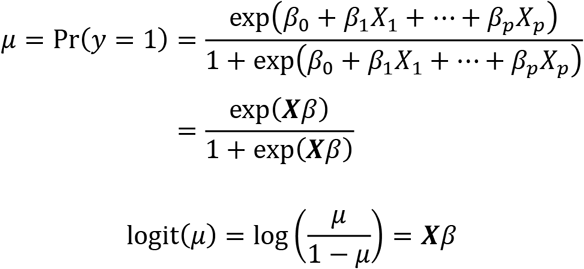

where *μ* = Pr(*y* = 1) denotes the probability that a genomic position is mutated (for testing individual SNV mutation types, e.g. A>T, *μ* is the probability that a site is mutated to a specific nucleotide), ***X*** represents the observations for the considered variables (categorical or continuous, e.g. ***d***_*mean*_, ***d***_*var*_, adjacent nucleotides, etc.), and *β* is the vector of parameters to be estimated.

We used the Bayesian logistic regression model implemented in the ‘bayesglm’ (Gelman et al. 2008) of the R package ‘arm’, which was reported to perform well in handling the complete separation issue in logistic regression models (Gelman et al. 2008). The complete separation issue is common when one class is rare relative to the other and (or) there are many regressors in a model. As we had only ~300,000 *de novo* mutations, the probability for a given site to be mutated in our data is ~1/10,000, which is a rare event.

Within the logistic regression framework, we compared the full model with all considered variables to a reduced model without one specific variable by performing likelihood-ratio tests in R (‘anova’ function) to evaluate the significance for each variable. The resulting p values of a set of likelihood-ratio tests were adjusted for multiple testing with Benjamini–Hochberg correction.

To perform the regression analysis, we generated the data of all variables for the *de novo* mutation sites and subsampled a fraction of the non-mutated sites as the control sites. We did not use all the non-mutated sites in the genome as it would lead to a large imbalance in the sizes of two classes (‘mutated’ and ‘non-mutated’) and much larger computational burden. For *de novo* SNVs, we randomly generated 2,561,953 non-mutated sites (about 1/1000 of the accessible genome, about 10 times as many as *de novo* SNVs) and 256,337 non-mutated sites (about 1/10,000 of the accessible genome, about 10 times as many as *de novo* INDELs) for INDELs. For *de novo* INDELs, we used the INDELs of ⩽5bp for regression analysis, because long INDELs were rare and may have high false positive/negative rates. For RNA expression, DNA methylation, chromatin accessibility, replication timing, recombination rate and histone modifications data, we used the average value of the ±10bp of a focal site for each specific feature based on the genome-wide signal tracks. We also assessed different window sizes (±5bp and ±20bp), which led to similar results.

For SNVs, we performed logistic regression tests for mutation types at A/T sites and C/G sites separately and distinguished C/G sites in CpG and non-CpG contexts. We also tested for nine individual SNV mutation types (three for A/T sites, three for C/G sites at CpG contexts, and three for non-CpG contexts, **Supplementary Fig. 3**). The regression coefficients for the full model of each test are given in **Supplementary Table 1.**

Since the variable ***d***_*mean*_ has a non-monotonic relationship with mutation rates, we binned the values into five categories: [0,18], [19, 36], [37, 54], [55, 73] and [74, 95] (first four bins implying nucleosome-bound regions, and the last bin implying close to the linker).

In the regression models mentioned above, we did not consider the non-additive effects of adjacent nucleotides (±5 bp). When we tried adding non-additive effects for ±5 nucleotides (considering only two-way interactions; taking a much longer running time), we got similar results regarding the association of translational stability (***d***_*var*_) and mutation rates (**Supplementary Fig. 4**). We also tried using the 7-mer mutability estimates from Carlson et al. (Carlson et al. 2018), which incorporated non-additive effects among ±3 nucleotides, as predictors in the regression models.

To evaluate how the sequence repeat status affects the effects of translational stability on mutation rates, We added the repeat status (‘Alu’, ‘L1’, ‘other repeat’ or ‘non-repeat’) as a predictor in the regression models, and also ran the regression tests for different repeat/non-repeat regions separately.

### Analysis of mutational processes

COSMIC mutational signatures are based on frequencies of mutations in tri-nucleotide contexts. Since the regions associated with strong nucleosomes have different tri-nucleotide composition relative to genome background, we first normalized the mutation type frequencies in regions associated with strong nucleosomes as this: set *F_i,strong_* for the occurrence of a specific mutation type (e,g. T[T>C]T), *N_i,strong_* for the occurrence of the considered tri-nucleotide context (e.g. TTT) in strong-nucleosome regions and *N_i,genome_* for the occurrence of the considered tri-nucleotide context in the whole-genome background, then the corrected occurrence of a the mutation type for strong nucleosomes is 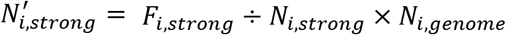. Fisher’s exact tests were performed to identify mutation types that show significant increase or decrease in strong-nucleosome regions relative to genome background. The contingency table used for running ‘fisher.test’ in R for a specific mutation type is *matrix*(*c*(*F_i,strong_, N_i,strong_ − F_i,strong_, F_i,genome_ − F_i,strong_*, (*N_i,genome_ − N_i,strong_*) − (*F_i,genome_ − F_i,strong_*)), *ncol* = 2), where *F_i,strong_*. And *F_i,genome_* are the occurrences of the considered mutation type and *N_i,strong_* and *N_i,genome_* for the occurrences of the considered tri-nucleotide context. Benjamini-Hochberg method was used for multiple testing correction.

The contribution of COSMIC mutational signatures (Alexandrov et al. 2013) to different sets of mutations (*de novo* SNVs and somatic mutations from bMMRD samples) was predicted using the ‘fit_to_signatures’ function in the R package ‘MutationalPatterns’ (Blokzijl et al. 2018). For the sets of *de novo* SNVs associated with strong nucleosomes, the corrected frequencies described above were used for running ‘fit_to_signatures’.

Mutations in *POLE* in cancers can lead to reduced base selectivity and/or deficient proofreading during replication, producing unusually large numbers of mutations (so called ‘ultra-hypermutation’) which facilitated our analysis. *POLE* mutated genomes from PCAWG project (Campbell et al. 2017) were used to evaluate the differential MMR efficiency between strong and non-strong nucleosome regions. We compared the mutation densities in cancer genomes with *POLE* mutated and a deficient MMR (4 individual samples) to those with *POLE* mutated and a proficient MMR (6 samples). The MMR pathway was considered deficient if a driver mutation (annotated by the PCAWG consortium) was found in one of five MMR core genes - *MLH1, MSH2, MSH6, PMS1* and *PMS2*.

For a given bin (10bp-size) in the meta-profile, we calculated the relative MMR escape ratio relative to genomic background around strong nucleosomes as described in the following formula,

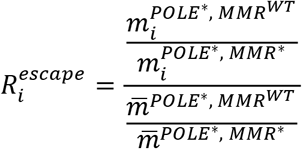

where *m_i_* is the mutation density for the *i*th bin (observed number of mutations in the *i*th bin divided by the bin size), and 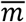 is the genome-wide average mutation density of a specific sample group (observed number of mutations in the simulated windows divided by the total window size), estimated by simulating random windows in the genome. A similar logic was used when evaluating relative proofreading escape ratios of Pol ε (mutated *POLE*) and Pol δ (mutated *POLD1*) using the somatic mutation data from the bMMRD project (Shlien et al. 2015).

When analyzing PCAWG and bMMRD data, to account for potential mappability issues, we focused on the highly mappable regions based on the CrgMapability scores from ENCODE. We used CrgMapability scores here, which are more stringent than GMA ones, because detecting somatic mutations in tumors is more difficult than for ordinary individual re-sequencing data. We considered the strong nucleosomes which have a 100mer CrgMapability score of 1 (meaning any 100-bp read from these regions can be mapped uniquely in the genome) within ±800bp of the dyads. We then simulated a same number of 1600bp-sized regions from the genome that satisfy the mappability requirement to calculate the background mutation density. Note that in theory the mappability issue in the relative escape ratios should be very small because the two sets of samples have the same mappability for a given bin and the ratio calculation normalizes the effects of different mappability among regions.

The raw reads of OK-seq data (Petryk et al. 2016) were downloaded from NCBI and mapped to the human genome. We kept only the uniquely mapped reads for inferring Okazaki junctions. The very 5’ end sites of aligned reads (separating reads mapped to Watson and Crick strands) were considered putative Okazaki junction signals.

To investigate DSBs around strong nucleosmes, we downloaded the genome-wide tracks of human END-seq data (GSM3227951 and GSM3227952) (Tubbs et al. 2018). Because the reads of END-seq data were single-ended 75bp, we considered the strong nucleosomes which have a 75mer CrgMapability score of 1 within ±500bp of the strong nucleosome dyads for analysis.

### Enrichment analysis for strong nucleosomes in different repeat contexts

GAT (Heger et al. 2013) was used to estimate the expected numbers of strong nucleosomes in different contexts (sampling >=1000 times), which were compared to the observed numbers. The annotations of repeat elements (Feb 2009, Repeat Library 20140131) were downloaded from RepeatMasker (Tempel 2012). We also did GAT analysis for LINE-1(L1) and Alu subfamilies of different ages. The age information of repeat families was from Giordano et al. (Giordano et al. 2007). For generating the MNase-seq midpoints along the repeat consensus sequences, we made use of the alignment information in the RepeatMasker result files (‘hg19.fa.align.gz’) and mapped the hg19-based coordinates to the coordinates in the consensus sequences. Strong nucleosomes appear to be under-detected in very young L1 elements, which we think is due to difficulties in mapping short MNase-seq reads (Alus are easier to map because they are much smaller).

Nucleosome deformation energies of all sites in the human genome were estimated using nuScore (Tolstorukov et al. 2008). We also used nuScore to estimate the deformation energies of Alu/L1 subfamily consensus sequences. For the L1 analysis shown in **Supplementary Fig. 8**, we only considered the 3’ end regions of L1 subfamilies, because 5’ end regions of L1 elements are usually truncated in the genome and their subfamily identities are difficult to be determined.

## Data availability

All the analyses in this study were based on published datasets.

## Acknowledgments

We are grateful to Tobias Warnecke, John Diffley, Anob Chakrabarti and Sara Rohban for insightful comments. We thank Peter Van Loo, Jonas Demeulemeester and Maxime Tarabichi for assistance in accessing the PCAWG genomic data. We appreciate obtaining access to the *de novo* mutation data on SFARI Base. This work is supported by the Francis Crick Institute which receives its core funding from Cancer Research UK (FC001110), the UK Medical Research Council (FC001110), and the Wellcome Trust (FC001110) (N.M.L.). N.M.L. is also supported by a Wellcome Trust Investigator Award and core funding from the Okinawa Institute of Science & Technology. C.L. is funded by an EMBO long-term postdoctoral fellowship (ALTF 1499-2016).

## Author contributions

C.L. conceived the project, performed the analyses and drafted the manuscript; N.M.L. supervised the project and co-wrote the manuscript.

## Supplementary Tables and Figures

**Supplementary Table 1 Coefficients of variables and other information from the full regression models for different mutation types (in a separate Excel file).** Note that for each of the categorical variables, the first category was used by the regression model as reference category (other categories were compared with the reference category) and thus there is no coefficient for that category. The statistics (4^th^ column) and p-values (5^th^ column) in the table were from Wald tests defaultly produced by ‘bayesglm’ (shown for reference), which are different from the likelihood ratio test-based p-values and were not used in our discussion.

**Supplementary Figure 1.**
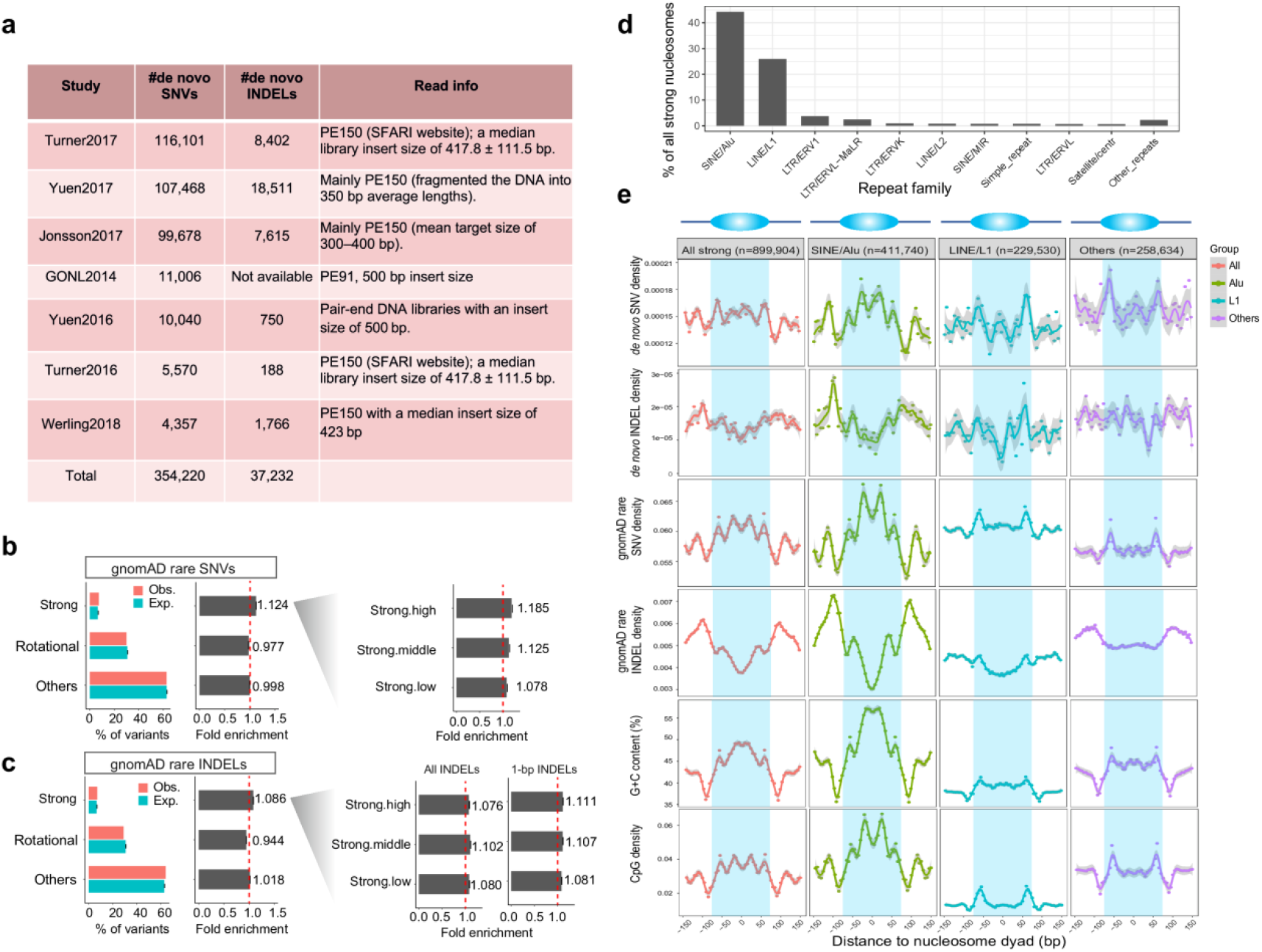
Mutations in different nucleosome contexts. (**a**) Information of the *de novo* mutation datasets from seven studies used in analysis. (**b**) Fold enrichment/depletion of gnomAD extremely rare SNVs in different nucleosome contexts. ‘Strong’, translationally stable positioning; ‘Rotational’, rotationally but not translationally stable positioning; ‘Others’, the remaining genomic regions. On the left is the fold enrichment for three subgroups of strong nucleosomes with different stabilities. Error bars depict 95% confidence intervals. (**c**) Fold enrichment/depletion of gnomAD INDELs in different nucleosome contexts. When using all INDELs the ‘strong.high’ group does not have a higher mutation rate than other two groups, but if using the 1-bp INDELs ‘strong.high’ does have the highest mutation rate among the three groups. We speculated that there may be more false negatives of longer INDELs in the ‘strong.high’ group. (**d**) Top 10 repeat families that are associated with strong nucleosomes. (**e**) Meta-profiles of SNV/INDEL densities (*de novo* or extremely rare variants) around all strong nucleosomes, or in different repeat-associated subgroups. At the bottom are the G+C content and CpG content profiles.

**Supplementary Figure 2.**
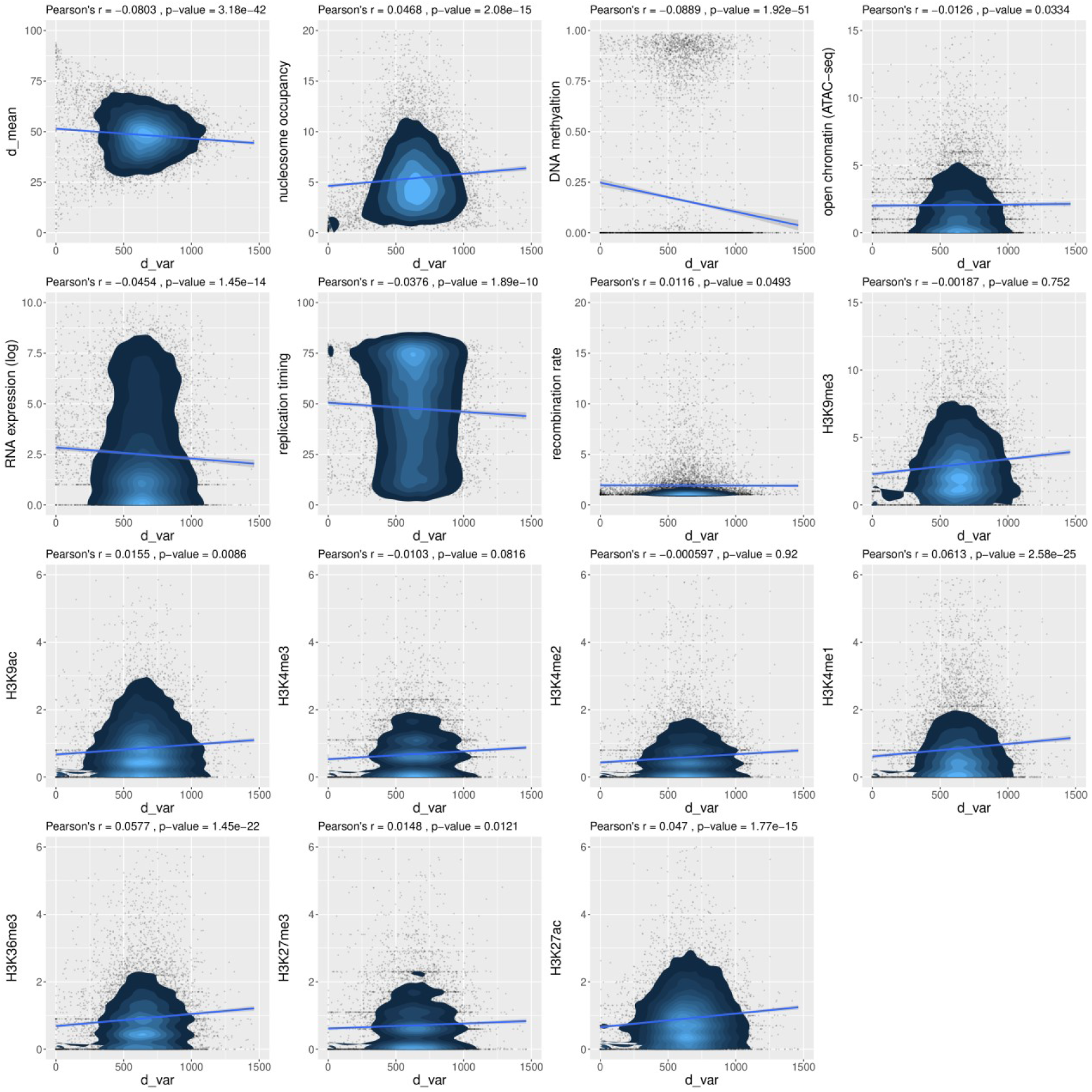
Correlation analysis between nucleosome positioning stability (*d_var_*) and other factors. On the top of each panel are the Pearson’s correlation coefficients and the corresponding p-values.

**Supplementary Figure 3.**
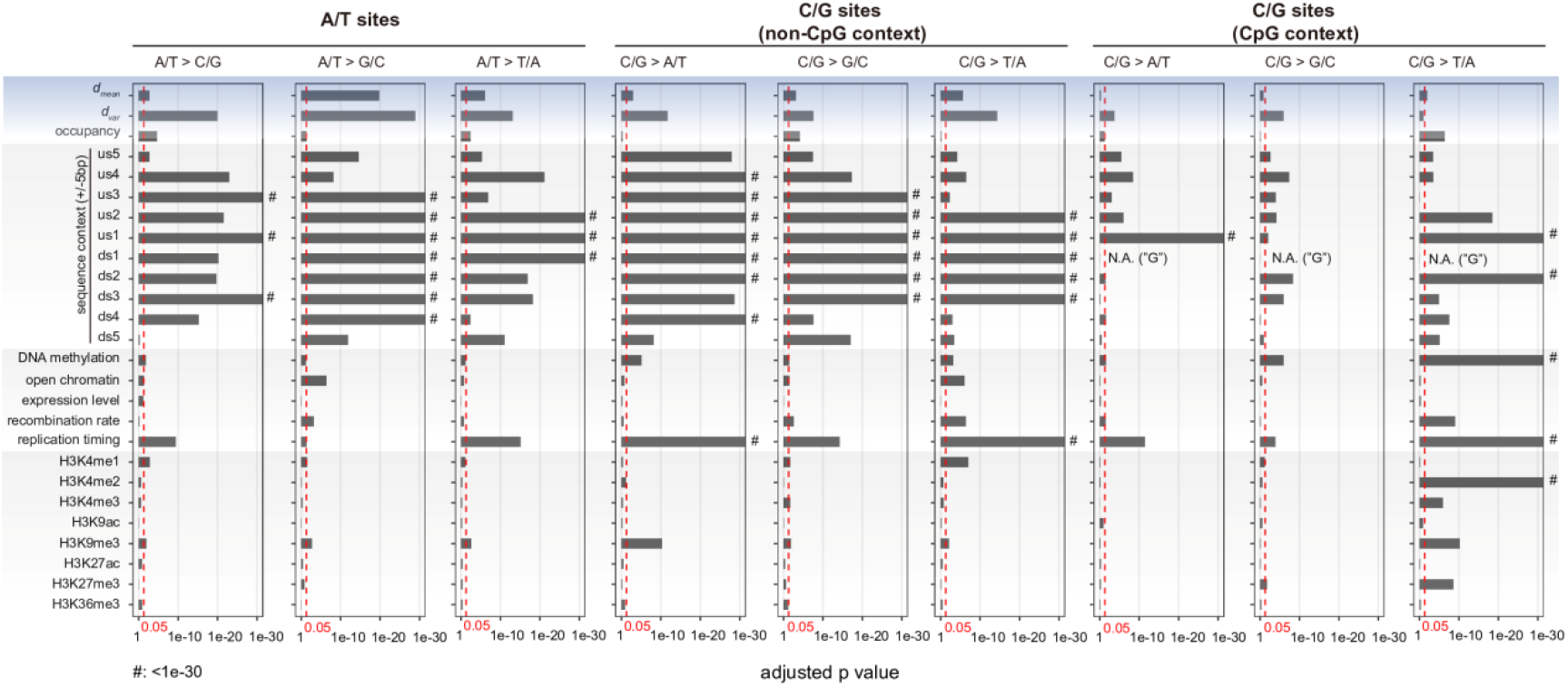
Results of statistical tests for nine individual SNV mutation types. C/G sites in non-CpG contexts and C/G sites in CpG contexts were tested separately. The red vertical lines represent the significance cut-off (0.05) for the adjusted p values (Benjamini–Hochberg correction). ‘us’, upstream; ‘ds’, downstream. ‘#’ means adjusted p < 1e-30.

**Supplementary Figure 4.**
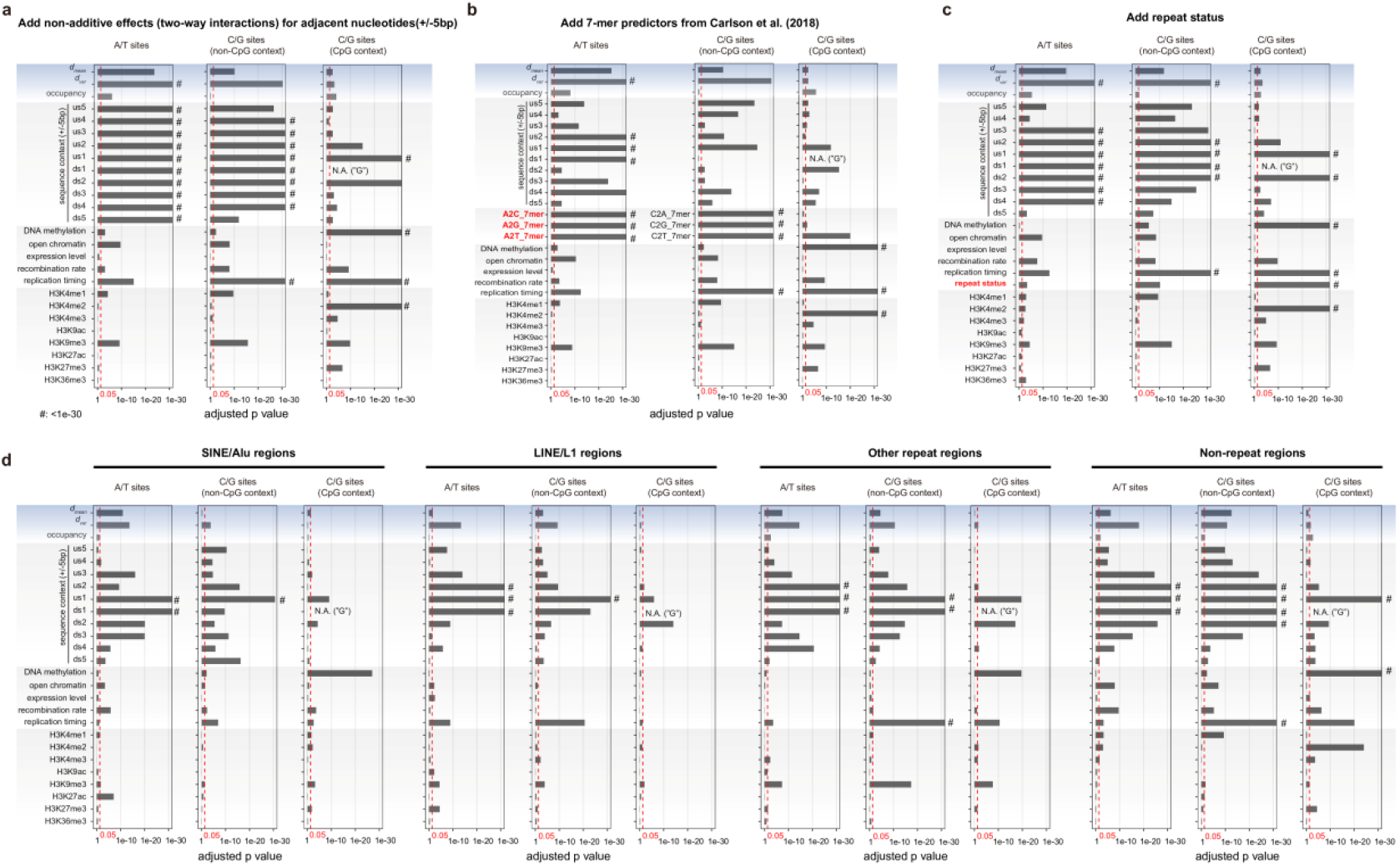
Results of statistical tests when considering two-way interactions of adjacent nucleotides, 7-mer mutability estimates from Carlson et al. and repeat status. (**a**) Adding the two-way interactions for ±5 nucleotides in the regression models. (**b**) Adding the 7-mer mutability estimates from Carlson et al. as predictors in the regression models. (**c**) Adding repeat status as a predictor in the regression models. (**d**) Running regression models for regions associated with different repeat contexts separately. We tested SNVs at A/T sites, C/G sites in non-CpG context and C/G sites in CpG context separately. The red vertical lines represent the significance cut-off (0.05) for the adjusted p values (Benjamini–Hochberg correction). ‘us’, upstream; ‘ds’, downstream. ‘#’ means adjusted p < 1e-30.

**Supplementary Figure 5.**
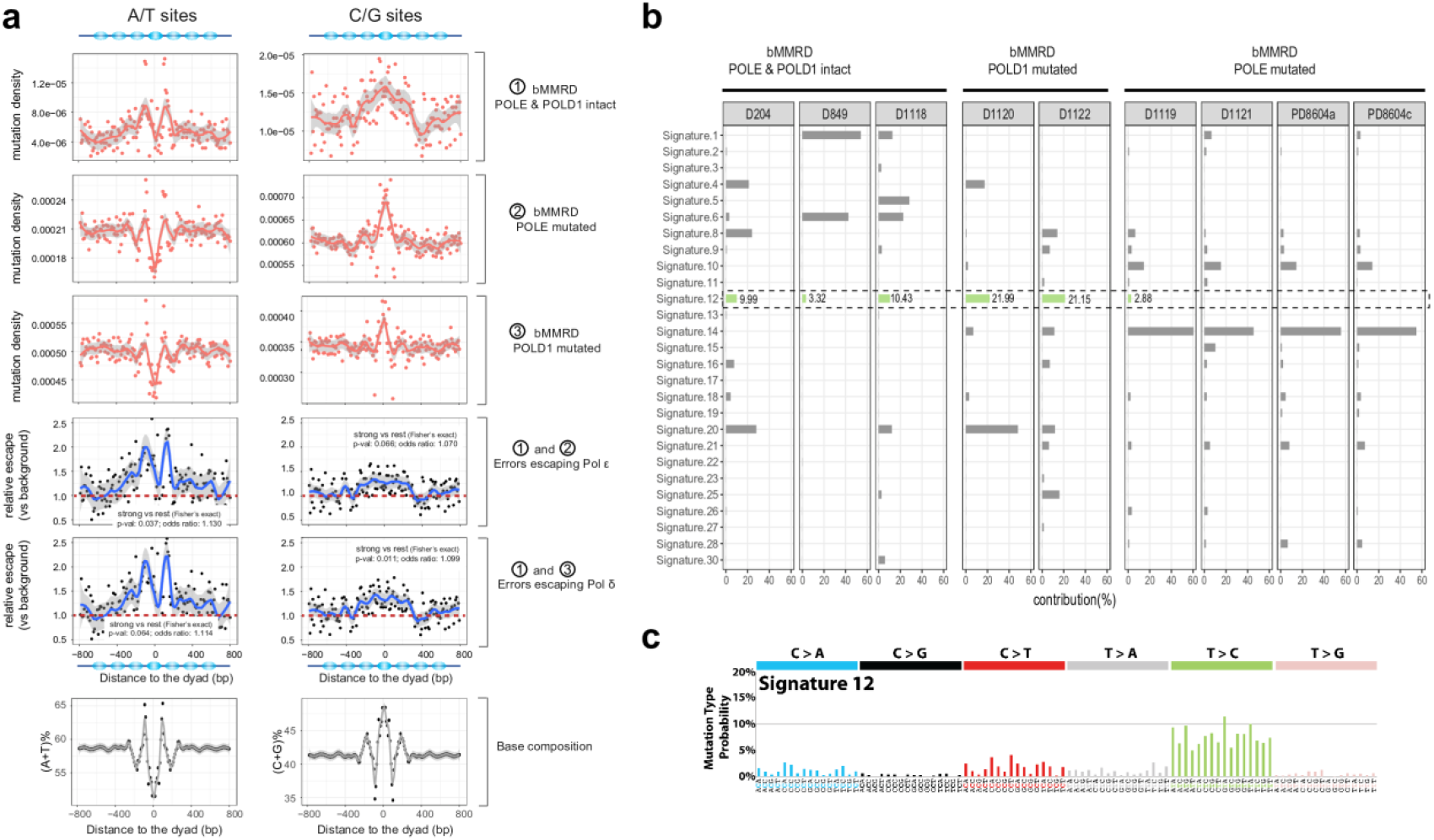
Analysis of related mutational processes using bMMRD data. (**a**) Mutation profiles around strong nucleosomes for bMMRD cancer genomes and the estimated relative escape ratios of Pol ε or Pol δ, for mutations at A/T sites and C/G sites respectively. Fisher’s exact test was used for testing the association of strong-nuclesome regions (dyad±95bp) with differential polymerase performance. (**b**) Comparison of the contribution of COSMIC mutational signatures predicted by MutationalPatterns in different bMMRD genomes. Highlighted is Signature 12, which shows a particularly high contribution in POLD1-muated bMMRD samples. (**c**) the tri-nucleotide mutational profile of Signature 12, obtained from COSMIC website.

**Supplementary Figure 6.**
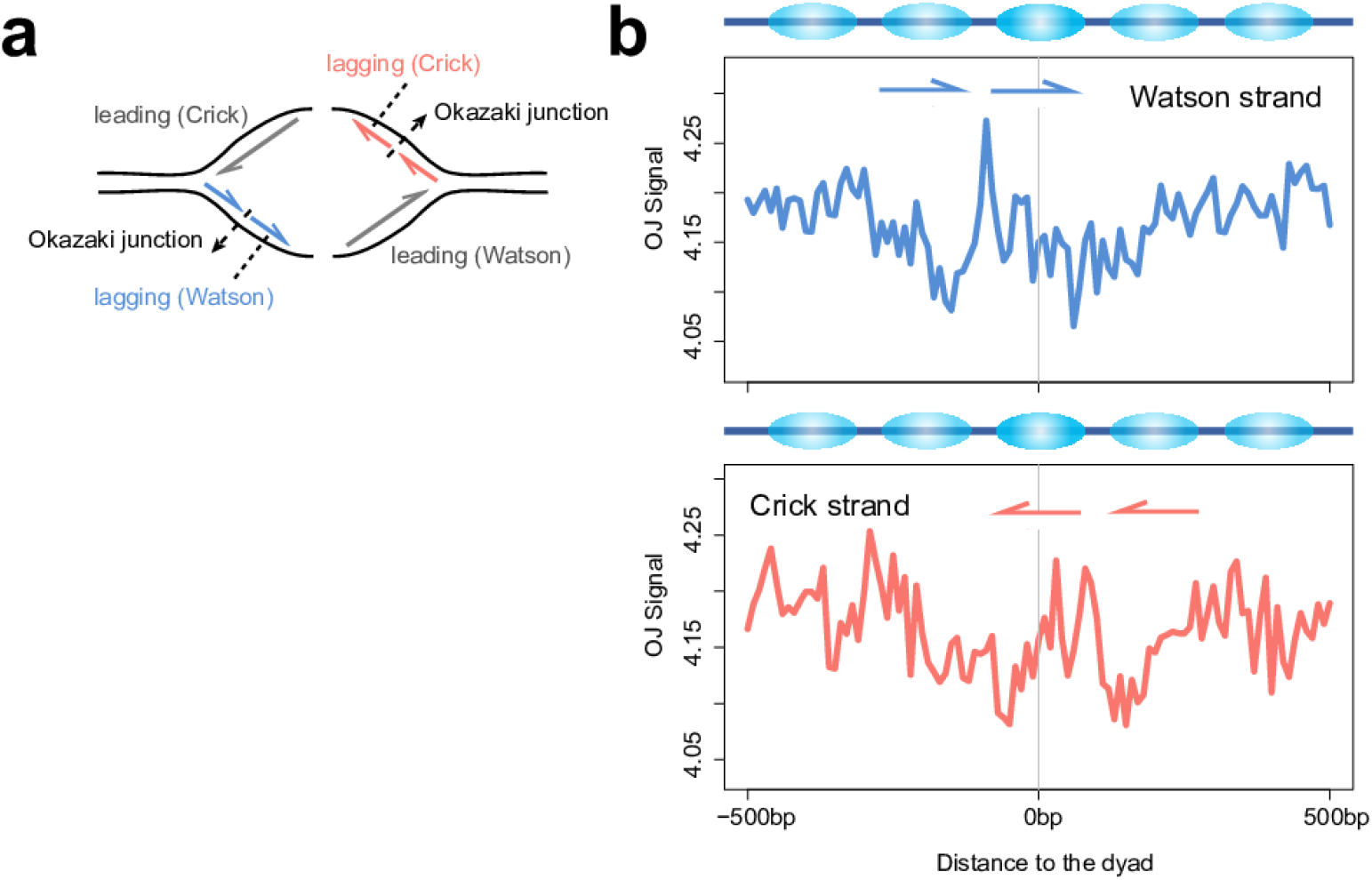
Analysis with OK-seq data. (**a**) Schematic illustrating replication strands and Okazaki junctions (OJs). (**b**) Meta-profile of the density of Okazaki junctions inferred from alignments of OK-seq reads around strong nucleosomes (high-mappability). OJ signals for Watson strand and Crick strand were plotted separately. Replication directions of Okazaki fragments are shown by arrows.

**Supplementary Figure 7.**
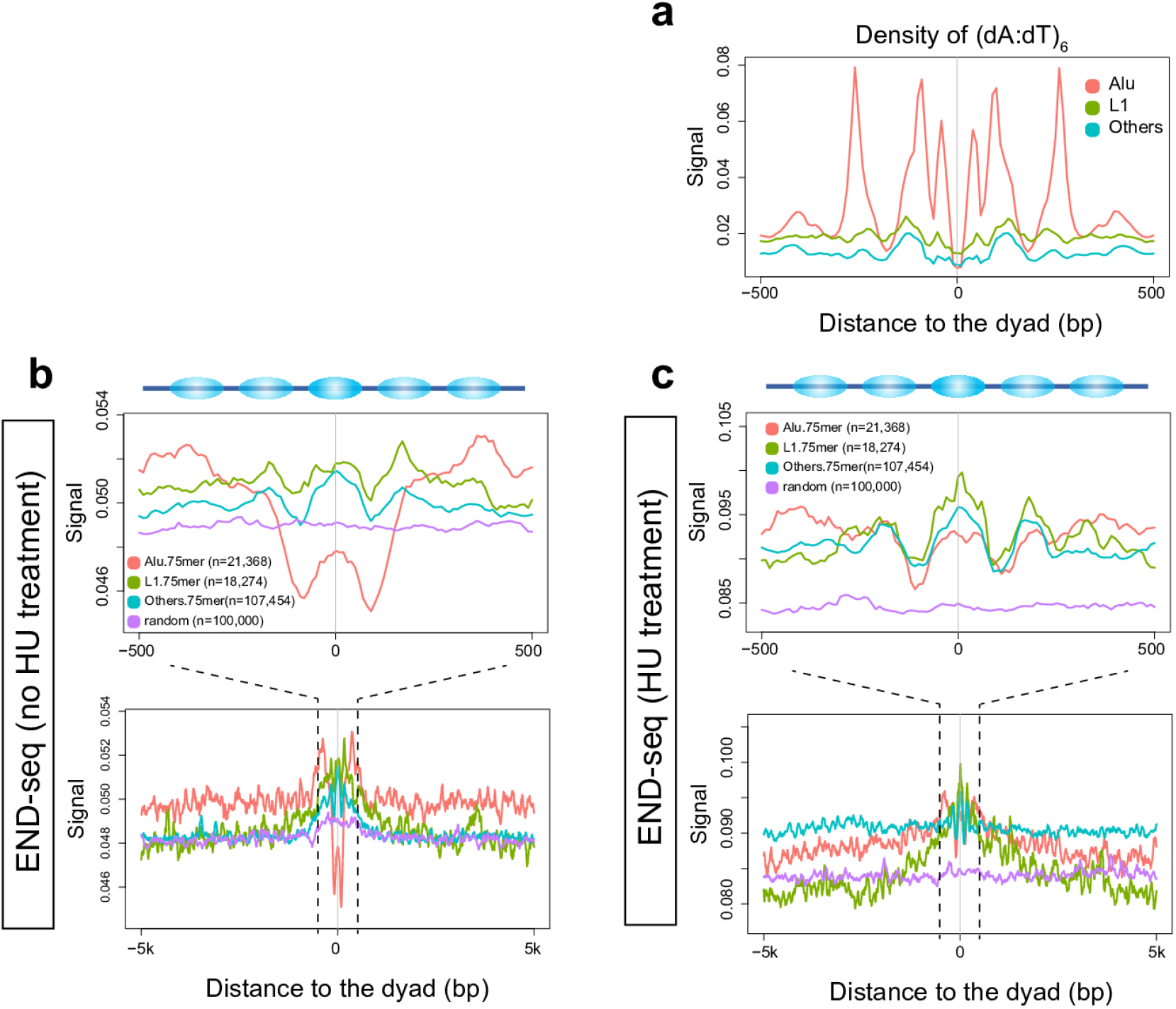
Analysis related to the DSBs around strong nucleosomes. (**a**) Density of poly(dA:dT) tracts (based on occurrence of (dA:dT)6 motifs) around strong nucleosomes. (**b-c**) Signal of DSBs based on the END-seq data around strong nucleosomes associated with different repeat elements. Only the strong nucleosomes of high 75-mer mappability within ±500bp were considered. Numbers of usable strong nucleosomes for each group are given in the brackets. HU (hydroxyurea) is a replicative stress-inducing agent.

**Supplementary Figure 8.**
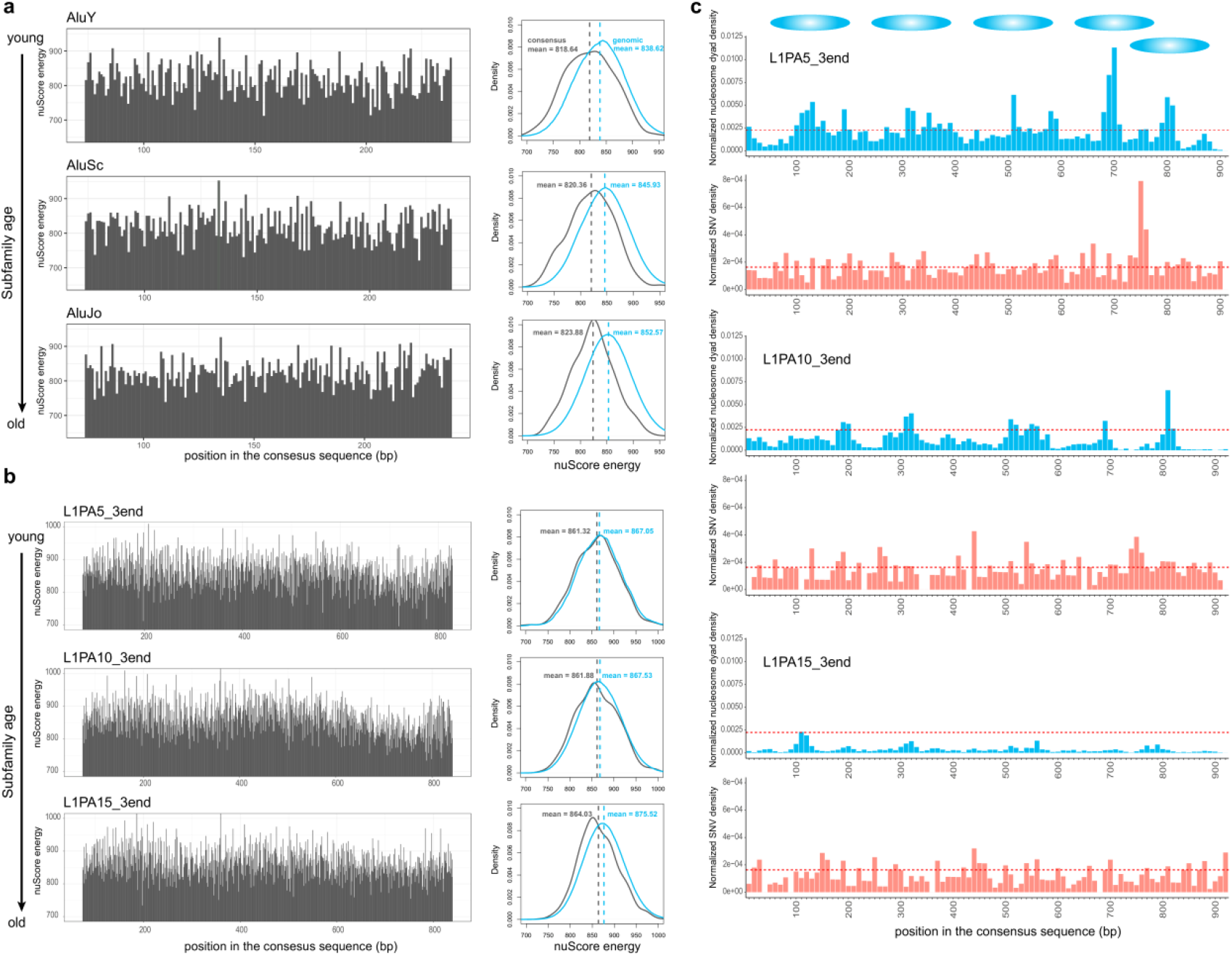
Additional analysis about repeat subfamily ages and strong nucleosomes. (**a**) nuScore-estimated per-base nucleosome deformation energies along three Alu subfamily consensus sequences. On the right are the comparisons of deformation energy distributions of the consensus sequences (ancestral states) and those of current genomic regions for the three subfamilies respectively. The deformation energy profiles of the consensus sequences are similar, but the average deformation energies increase over time, with older Alu subfamilies displaying larger differences relative to the consensus. (**b**) Similar to (a), but for three example L1 subfamilies. (**c**) Barplots for normalized densities of strong nucleosome dyads and *de novo* SNVs along the consensus sequences of three L1 subfamilies, using 10-bp bins. Several loci that are enriched for dyads of strong nucleosomes are shown on the top with ellipses. The red dash lines represent the average densities for the L1PA5 subfamily. The densities of strong nucleosome dyads and *de novo* SNVs appear to decrease over evolutionary time.

## References

Acuna-Hidalgo R, Veltman JA, Hoischen A. 2016. New insights into the generation and role of de novo mutations in health and disease. Genome biology 17(1): 241.

Alexandrov LB, Nik-Zainal S, Wedge DC, Aparicio SA, Behjati S, Biankin AV, Bignell GR, Bolli N, Borg A, Borresen-Dale AL et al. 2013. Signatures of mutational processes in human cancer. Nature 500(7463): 415–421.

Blokzijl F, Janssen R, van Boxtel R, Cuppen E. 2018. MutationalPatterns: comprehensive genome-wide analysis of mutational processes. Genome medicine 10(1): 33.

Campbell PJ, Getz G, Stuart JM, Korbel JO, Stein LD. 2017. Pan-cancer analysis of whole genomes. BioRxiv: 162784.

Carlson J, Locke AE, Flickinger M, Zawistowski M, Levy S, Myers RM, Boehnke M, Kang HM, Scott LJ, Li JZ et al. 2018. Extremely rare variants reveal patterns of germline mutation rate heterogeneity in humans. Nature communications 9(1): 3753.

Chen X, Chen Z, Chen H, Su Z, Yang J, Lin F, Shi S, He X. 2012. Nucleosomes suppress spontaneous mutations base-specifically in eukaryotes. Science 335(6073): 1235–1238.

Drillon G, Audit B, Argoul F, Arneodo A. 2016. Evidence of selection for an accessible nucleosomal array in human. BMC genomics 17: 526.

ENCODE Consortium. 2012. An integrated encyclopedia of DNA elements in the human genome. Nature 489(7414): 57–74.

Englander EW, Howard BH. 1995. Nucleosome positioning by human Alu elements in chromatin. The Journal of biological chemistry 270(17): 10091–10096.

Englander EW, Wolffe AP, Howard BH. 1993. Nucleosome interactions with a human Alu element. Transcriptional repression and effects of template methylation. The Journal of biological chemistry 268(26): 19565–19573.

Francioli LC, Polak PP, Koren A, Menelaou A, Chun S, Renkens I, Genome of the Netherlands C, van Duijn CM, Swertz M, Wijmenga C et al. 2015. Genome-wide patterns and properties of de novo mutations in humans. Nature genetics 47(7): 822–826.

Frigola J, Sabarinathan R, Mularoni L, Muinos F, Gonzalez-Perez A, Lopez-Bigas N. 2017. Reduced mutation rate in exons due to differential mismatch repair. Nature genetics 49(12): 1684–1692.

Gaffney DJ, McVicker G, Pai AA, Fondufe-Mittendorf YN, Lewellen N, Michelini K, Widom J, Gilad Y, Pritchard JK. 2012. Controls of nucleosome positioning in the human genome. PLoS genetics 8(11): e1003036.

Gelman A, Jakulin A, Pittau MG, Su Y-S. 2008. A weakly informative default prior distribution for logistic and other regression models. The Annals of Applied Statistics 2(4): 1360–1383.

Genome of the Netherlands C. 2014. Whole-genome sequence variation, population structure and demographic history of the Dutch population. Nature genetics 46(8): 818–825.

Giordano J, Ge Y, Gelfand Y, Abrusan G, Benson G, Warburton PE. 2007. Evolutionary history of mammalian transposons determined by genome-wide defragmentation. PLoS computational biology 3(7): e137.

Guo J, Grow EJ, Yi C, Mlcochova H, Maher GJ, Lindskog C, Murphy PJ, Wike CL, Carrell DT, Goriely A et al. 2017. Chromatin and Single-Cell RNA-Seq Profiling Reveal Dynamic Signaling and Metabolic Transitions during Human Spermatogonial Stem Cell Development. Cell stem cell 21(4): 533–546 e536.

Harpak A, Bhaskar A, Pritchard JK. 2016. Mutation Rate Variation is a Primary Determinant of the Distribution of Allele Frequencies in Humans. PLoS genetics 12(12): e1006489.

Harris K, Nielsen R. 2014. Error-prone polymerase activity causes multinucleotide mutations in humans. Genome research 24(9): 1445–1454.

Heger A, Webber C, Goodson M, Ponting CP, Lunter G. 2013. GAT: a simulation framework for testing the association of genomic intervals. Bioinformatics 29(16): 2046–2048.

Hodgkinson A, Eyre-Walker A. 2011. Variation in the mutation rate across mammalian genomes. Nature reviews Genetics 12(11): 756–766.

International HapMap Consortium Frazer KA Ballinger DG Cox DR Hinds DA Stuve LL Gibbs RA Belmont JW Boudreau A Hardenbol P et al. 2007. A second generation human haplotype map of over 3.1 million SNPs. Nature 449(7164): 851–861.

Jonsson H, Sulem P, Kehr B, Kristmundsdottir S, Zink F, Hjartarson E, Hardarson MT, Hjorleifsson KE, Eggertsson HP, Gudjonsson SA et al. 2017. Parental influence on human germline de novo mutations in 1,548 trios from Iceland. Nature 549(7673): 519–522.

Lee H, Schatz MC. 2012. Genomic dark matter: the reliability of short read mapping illustrated by the genome mappability score. Bioinformatics 28(16): 2097–2105.

Lek M, Karczewski KJ, Minikel EV, Samocha KE, Banks E, Fennell T, O’Donnell-Luria AH, Ware JS, Hill AJ, Cummings BB et al. 2016. Analysis of protein-coding genetic variation in 60,706 humans. Nature 536(7616): 285–291.

Li F, Tian L, Gu L, Li GM. 2009. Evidence that nucleosomes inhibit mismatch repair in eukaryotic cells. The Journal of biological chemistry 284(48): 33056–33061.

Lujan SA, Clausen AR, Clark AB, MacAlpine HK, MacAlpine DM, Malc EP, Mieczkowski PA, Burkholder AB, Fargo DC, Gordenin DA et al. 2014. Heterogeneous polymerase fidelity and mismatch repair bias genome variation and composition. Genome research 24(11): 1751–1764.

Makova KD, Hardison RC. 2015. The effects of chromatin organization on variation in mutation rates in the genome. Nature reviews Genetics 16(4): 213–223.

Michaelson JJ, Shi Y, Gujral M, Zheng H, Malhotra D, Jin X, Jian M, Liu G, Greer D, Bhandari A et al. 2012. Whole-genome sequencing in autism identifies hot spots for de novo germline mutation. Cell 151(7): 1431–1442.

Perera D, Poulos RC, Shah A, Beck D, Pimanda JE, Wong JW. 2016. Differential DNA repair underlies mutation hotspots at active promoters in cancer genomes. Nature 532(7598): 259–263.

Petryk N, Kahli M, d’Aubenton-Carafa Y, Jaszczyszyn Y, Shen Y, Silvain M, Thermes C, Chen CL, Hyrien O. 2016. Replication landscape of the human genome. Nature communications 7: 10208.

Pich O, Muinos F, Sabarinathan R, Reyes-Salazar I, Gonzalez-Perez A, Lopez-Bigas N. 2018. Somatic and Germline Mutation Periodicity Follow the Orientation of the DNA Minor Groove around Nucleosomes. Cell 175(4): 1074–1087 e1018.

Prendergast JG, Semple CA. 2011. Widespread signatures of recent selection linked to nucleosome positioning in the human lineage. Genome research 21(11): 1777–1787.

Ramirez F, Dundar F, Diehl S, Gruning BA, Manke T. 2014. deepTools: a flexible platform for exploring deep-sequencing data. Nucleic acids research 42(Web Server issue): W187–191.

Reijns MAM, Kemp H, Ding J, de Proce SM, Jackson AP, Taylor MS. 2015. Lagging-strand replication shapes the mutational landscape of the genome. Nature 518(7540): 502–506.

Rodgers K, McVey M. 2016. Error-Prone Repair of DNA Double-Strand Breaks. Journal of cellular physiology 231(1): 15–24.

Roy S, Tomaszowski KH, Luzwick JW, Park S, Li J, Murphy M, Schlacher K. 2018. p53 orchestrates DNA replication restart homeostasis by suppressing mutagenic RAD52 and POLtheta pathways. eLife 7.

Sabarinathan R, Mularoni L, Deu-Pons J, Gonzalez-Perez A, Lopez-Bigas N. 2016. Nucleotide excision repair is impaired by binding of transcription factors to DNA. Nature 532(7598): 264–267.

Salih F, Salih B, Kogan S, Trifonov EN. 2008. Epigenetic nucleosomes: Alu sequences and CG as nucleosome positioning element. Journal of biomolecular structure & dynamics 26(1): 9–16.

Sasaki S, Mello CC, Shimada A, Nakatani Y, Hashimoto S, Ogawa M, Matsushima K, Gu SG, Kasahara M, Ahsan B et al. 2009. Chromatin-associated periodicity in genetic variation downstream of transcriptional start sites. Science 323(5912): 401–404.

Schuster-Bockler B, Lehner B. 2012. Chromatin organization is a major influence on regional mutation rates in human cancer cells. Nature 488(7412): 504–507.

Segurel L, Wyman MJ, Przeworski M. 2014. Determinants of mutation rate variation in the human germline. Annual review of genomics and human genetics 15: 47–70.

Seplyarskiy VB, Akkuratov EE, Akkuratova N, Andrianova MA, Nikolaev SI, Bazykin GA, Adameyko I, Sunyaev SR. 2018. Error-prone bypass of DNA lesions during lagging-strand replication is a common source of germline and cancer mutations. Nature genetics.

Seplyarskiy VB, Andrianova MA, Bazykin GA. 2017. APOBEC3A/B-induced mutagenesis is responsible for 20% of heritable mutations in the TpCpW context. Genome research 27(2): 175–184.

Shlien A, Campbell BB, de Borja R, Alexandrov LB, Merico D, Wedge D, Van Loo P, Tarpey PS, Coupland P, Behjati S et al. 2015. Combined hereditary and somatic mutations of replication error repair genes result in rapid onset of ultra-hypermutated cancers. Nature genetics 47(3): 257–262.

Slotkin RK, Martienssen R. 2007. Transposable elements and the epigenetic regulation of the genome. Nature reviews Genetics 8(4): 272–285.

Smerdon MJ. 1991. DNA repair and the role of chromatin structure. Current opinion in cell biology 3(3): 422–428.

Smith TCA, Arndt PF, Eyre-Walker A. 2018. Large scale variation in the rate of germ-line de novo mutation, base composition, divergence and diversity in humans. PLoS genetics 14(3): e1007254.

Stamatoyannopoulos JA, Adzhubei I, Thurman RE, Kryukov GV, Mirkin SM, Sunyaev SR. 2009. Human mutation rate associated with DNA replication timing. Nature genetics 41(4): 393–395.

Supek F, Lehner B. 2015. Differential DNA mismatch repair underlies mutation rate variation across the human genome. Nature 521(7550): 81–84.

Tanaka Y, Yamashita R, Suzuki Y, Nakai K. 2010. Effects of Alu elements on global nucleosome positioning in the human genome. BMC genomics 11: 309.

Tempel S. 2012. Using and understanding RepeatMasker. Methods in molecular biology 859: 29–51.

Terekhanova NV, Seplyarskiy VB, Soldatov RA, Bazykin GA. 2017. Evolution of Local Mutation Rate and Its Determinants. Molecular biology and evolution 34(5): 1100–1109.

Tolstorukov MY, Choudhary V, Olson WK, Zhurkin VB, Park PJ. 2008. nuScore: a web-interface for nucleosome positioning predictions. Bioinformatics 24(12): 1456–1458.

Tolstorukov MY, Volfovsky N, Stephens RM, Park PJ. 2011. Impact of chromatin structure on sequence variability in the human genome. Nature structural & molecular biology 18(4): 510–515.

Tomasetti C, Li L, Vogelstein B. 2017. Stem cell divisions, somatic mutations, cancer etiology, and cancer prevention. Science 355(6331): 1330–1334.

Tomasetti C, Vogelstein B. 2015. Variation in cancer risk among tissues can be explained by the number of stem cell divisions. Science 347(6217): 78–81.

Tubbs A, Sridharan S, van Wietmarschen N, Maman Y, Callen E, Stanlie A, Wu W, Wu X, Day A, Wong N et al. 2018. Dual Roles of Poly(dA:dT) Tracts in Replication Initiation and Fork Collapse. Cell 174(5): 1127–1142 e1119.

Turner TN, Coe BP, Dickel DE, Hoekzema K, Nelson BJ, Zody MC, Kronenberg ZN, Hormozdiari F, Raja A, Pennacchio LA et al. 2017a. Genomic Patterns of De Novo Mutation in Simplex Autism. Cell 171(3): 710–722 e712.

Turner TN, Hormozdiari F, Duyzend MH, McClymont SA, Hook PW, Iossifov I, Raja A, Baker C, Hoekzema K, Stessman HA et al. 2016. Genome Sequencing of Autism-Affected Families Reveals Disruption of Putative Noncoding Regulatory DNA. American journal of human genetics 98(1): 58–74.

Turner TN, Yi Q, Krumm N, Huddleston J, Hoekzema K, HA FS, Doebley AL, Bernier RA, Nickerson DA, Eichler EE. 2017b. denovo-db: a compendium of human de novo variants. Nucleic acids research 45(D1): D804–D811.

Veltman JA, Brunner HG. 2012. De novo mutations in human genetic disease. Nature reviews Genetics 13(8): 565–575.

Warnecke T, Becker EA, Facciotti MT, Nislow C, Lehner B. 2013. Conserved substitution patterns around nucleosome footprints in eukaryotes and Archaea derive from frequent nucleosome repositioning through evolution. PLoS computational biology 9(11): e1003373.

Werling DM, Brand H, An JY, Stone MR, Zhu L, Glessner JT, Collins RL, Dong S, Layer RM, Markenscoff-Papadimitriou E et al. 2018. An analytical framework for whole-genome sequence association studies and its implications for autism spectrum disorder. Nature genetics 50(5): 727–736.

West JA, Cook A, Alver BH, Stadtfeld M, Deaton AM, Hochedlinger K, Park PJ, Tolstorukov MY, Kingston RE. 2014. Nucleosomal occupancy changes locally over key regulatory regions during cell differentiation and reprogramming. Nature communications 5: 4719.

Winter DR, Song L, Mukherjee S, Furey TS, Crawford GE. 2013. DNase-seq predicts regions of rotational nucleosome stability across diverse human cell types. Genome research 23(7): 1118–1129.

Yuen R, Merico D, Bookman M, J LH, Thiruvahindrapuram B, Patel RV, Whitney J, Deflaux N, Bingham J, Wang Z et al. 2017. Whole genome sequencing resource identifies 18 new candidate genes for autism spectrum disorder. Nature neuroscience 20(4): 602–611.

Yuen RK, Merico D, Cao H, Pellecchia G, Alipanahi B, Thiruvahindrapuram B, Tong X, Sun Y, Cao D, Zhang T et al. 2016. Genome-wide characteristics of de novo mutations in autism. NPJ genomic medicine 1: 160271–1602710.

